# SpliceMutr enables pan-cancer analysis of splicing-derived neoantigen burden in tumors

**DOI:** 10.1101/2023.05.26.542165

**Authors:** Theron Palmer, Michael D Kessler, Xiaoshan M. Shao, Archana Balan, Mark Yarchoan, Neeha Zaidi, Tamara Y Lopez-Vidal, Ali Saeed, Jessica Gore, Nilofer S Azad, Elizabeth M Jaffee, Alexander V Favorov, Valsamo Anagnostou, Rachel Karchin, Daria A Gaykalova, Ludmila Danilova, Elana J Fertig

## Abstract

Aberrant alternative splicing can generate neoantigens, which can themselves stimulate immune responses and surveillance. Previous methods for quantifying splicing-derived neoantigens are limited by independent references and potential batch effects. Here, we introduce SpliceMutr, a bioinformatics approach and pipeline for identifying splicing derived neoantigens from tumor and normal data. SpliceMutr facilitates the identification of tumor-specific antigenic splice variants, predicts MHC-binding affinity, and estimates splicing antigenicity scores per gene. By applying this tool to genomic data from The Cancer Genome Atlas (TCGA), we generate splicing-derived neoantigens and neoantigenicity scores per sample and across all cancer types and find numerous correlations between splicing antigenicity and well-established biomarkers of anti-tumor immunity. Notably, carriers of mutations within splicing machinery genes have higher splicing antigenicity, which provides support for our approach. Further analysis of splicing antigenicity in cohorts of melanoma patients treated with mono-or combined immune checkpoint inhibition suggest that the abundance of splicing antigens is reduced post-treatment from baseline in patients who progress, likely because of an immunoediting process. We also observe increased splicing antigenicity in responders to immunotherapy, which may relate to an increased capacity to mount an immune response to splicing-derived antigens. We find the splicing antigenicity to be higher in tumor samples when compared to normal, that mutations in the splicing machinery result in increased splicing antigenicity in some cancers, and higher splicing antigenicity is associated with positive response to immune checkpoint inhibitor therapies. Further, this new computational pipeline provides novel analytical capabilities for splicing antigenicity and is openly available for further immuno-oncologic analysis.

## Background

Immune surveillance of premalignant or cancerous lesions is contingent on the recognition of tumor-specific neoantigens presented on the surface of tumor cells and their elimination by the immune system. Despite the continuous pressure by the immune system to prevent the growth of nascent tumor clones, cancer cells evolve mechanisms of immune suppression that prevent their effective elimination and enable their escape and growth. Upregulation of immune checkpoints, such as CTLA-4, by regulatory T cells, Tregs, or PD-1/PD-L1 axis in the tumor microenvironment results in abrogation of effector T cell function that compromises immune clearance of tumor clones. This sets the basis for the clinical development of immune checkpoint inhibitors (ICI) that remove the brakes of these inhibitory checkpoints and reinvigorate antitumor immunity (Hong & Maleki Vareki, 2022; Wu et al., 2022). Identifying biomarkers and functional determinants of response to ICI is an active area of research.

Tumor mutational burden (TMB) has been widely recognized as a biomarker of response to immunotherapy, as a high mutational burden is thought to give rise to more neoantigens that can be recognized and eliminated by immune cells. However, TMB is not a universal biomarker of immune checkpoint inhibition (ICI) response or cytotoxic T-cell infiltration (Wu et al., 2022). While mutations represent one source of alteration to the protein structure of genes in cancer, further transcriptional and translational modifications to protein structure can also induce tumor antigens independent of mutation status if these modifications introduce altered peptide sequences to the protein. Identifying these mutation-independent tumor antigens is key to mapping the determinants of immunologic tumor control and subsequent clinical responses.

Currently, several computational algorithms predict immunogenic neoantigens based on the extraction of various features, including expression levels of mutant alleles, MHC presentation, mutant epitope-MHC binding affinity, and probability of being recognized by the T cells. These algorithms focused on tumor-specific non-synonymous variants, including indels. However, given the incomplete ability of TMB and canonical predicted neoantigens to accurately stratify patients based on response to ICI therapies, alternate sources of neoantigen production and methodologies to identify them are needed.

Abnormal RNA splicing in cancer cells contributes to many oncogenic processes, and splicing modulators have emerged as attractive targets for cancer therapy (Inoue et al., 2019; Xu et al., 2019; Haen et al., 2020; Liu et al., 2020a; Jayasinghe et al., 2018a). The association between splicing dysregulation and immune recognition across cancers has been explored in previous analyses (Jayasinghe et al., 2018; Kahles et al., 2018). Notably, previous work showed that in breast and ovarian cancers, the number of MHC-bound peptides derived from tumor-specific splicing events was higher, on average, than those resulting from single nucleotide variants (Kahles et al. 2018). The capacity of the splicing events to create splicing-impacted peptides that can bind to the MHC complex suggests that they may have translational implications on the outcome of immunotherapy treatment. For example, a study evaluating the anticancer drug indisulam, an RBM39 inhibitor that disrupts RNA splicing, in a mouse model of melanoma found that MHC-I presentation of splicing-derived immunogenic peptides induced T cell activation and amplified the therapeutic efficacy of PD-1 inhibition (Lu et al., 2021). Despite mounting evidence that dysregulated splicing represents a source of viable tumor neoantigens, their functional relevance and clinical significance in the context of immunotherapy responses remain poorly explored and require in-depth studies.

Most current computational pipelines for neoantigen prediction do not consider splicing-derived neoantigens and thus overlook a potentially large fraction of the neoantigen repertoire that may be informative of antitumor immune responses. Moreover, current methodologies for splicing neoantigen analysis using short-read RNA-seq data depend upon splice-graph augmentation (Kahles et al., 2016, 2018) or intron-centric outlier analysis using mixed-tissue references (Trincado et al., 2021). Further, they do not leverage intron-centric differential or outlier splicing tools like LeafCutter (Y. I. Li et al., 2018), SEVA (Afsari et al., 2018), or LeafCutterMD (Jenkinson et al., 2020). Splice graph augmentation by short-read RNA-seq data is carried out by generating a splicing graph based on a reference transcriptome and then augmenting that reference to represent the transcriptome evidenced by RNA-seq reads. Similar to splice-graph assembly, splice-graph augmentation can result in fragmented transcriptomes. Intron-centric outlier analysis can be carried out by comparing identified introns from target and mixed tissue references and filtering out any target introns that are also included in the mixed tissue reference (Trincado et al., 2021). Intron-centric outlier analysis can introduce unwanted batch effects when using mixed-tissue references. Intron-centric differential (Afsari et al., 2018; Y. I. Li et al., 2018) or outlier (Jenkinson et al., 2020) splicing analysis leverages the strengths of short-read sequencing, primarily high accuracy, against its main weakness, non-multi-exon spanning reads. Still, further extensions to these rigorous methods for short-read differential or outlier splicing analysis methods are also needed to fully address the potential of the predicted splice forms to create protein-coding products and provide a quantitative means to comparing splicing antigenicity to other molecular biomarkers used to evaluate ICI response.

In the current study, we have developed a methodology based on intron-centric splicing analysis, SpliceMutr [https://github.com/FertigLab/splicemute.git], that calculates tumor and normal aberrant splicing-derived neoantigen load per gene and per sample that we use to compare splicing antigenicity to current measures of the immune response. We further apply SpliceMutr to the TCGA dataset to define the interplay between the splicing antigenicity and TMB. We also checked the role mutations in the splicing factor machinery play in the level of splicing antigenicity. Finally, we used a cohort of ICI-treated melanoma patients to define how splicing antigenicity relates to clinical outcomes with immune checkpoint blockade. Altogether, our analyses with SpliceMutr lead us to hypothesize that the diversity of neoantigens resulting from the tumor splicing antigenicity increases the ability of the immune system to mount a successful immune response, which warrants future mechanistic studies.

## Implementation

### Overview of the SpliceMutr pipeline for differential splicing antigenicity analysis

SpliceMutr is a comprehensive computational pipeline that identifies splicing-derived neoantigens through differential analysis of short-read RNA-sequencing data between two given groups of samples, illustrated for tumor and normal samples in Fig 1. Due to replicates being required for the differential splicing analysis performed using the SpliceMutr pipeline, the tumor, and normal sample inputs are unmatched in this study and should be for future applications. This pipeline leverages both R and Python scripts to enable a new analysis approach for the identification of candidate neoepitopes. SpliceMutr relies on aligned RNA-seq reads using a splicing-aware aligner, such as STAR, and the class 1 HLA genotype per sample, which can be computed from the RNA-seq data (Orenbuch et al., 2020). The split-read genomic coordinates and counts are then processed and used to perform differential or outlier intron-usage analysis using LeafCutter (Y. I. Li et al., 2018). SpliceMutr uses the results of this differential intron usage analysis as the basis of transcript formation, ORF **L**ength and **GC** content (LGC) coding potential prediction (Wang et al., 2019), translation, and peptide kmerization. Additionally, SpliceMutr uses the LeafCutter-identified differential intron clusters and their associated SpliceMutr-formed peptides to filter out kmer sequences shared between two groups of interest. Using the per-sample HLA-genotype and the kmers associated with each transcript, SpliceMutr runs MHCnuggets (Shao et al., 2020) to predict MHC binding affinity.

**Fig 1.**
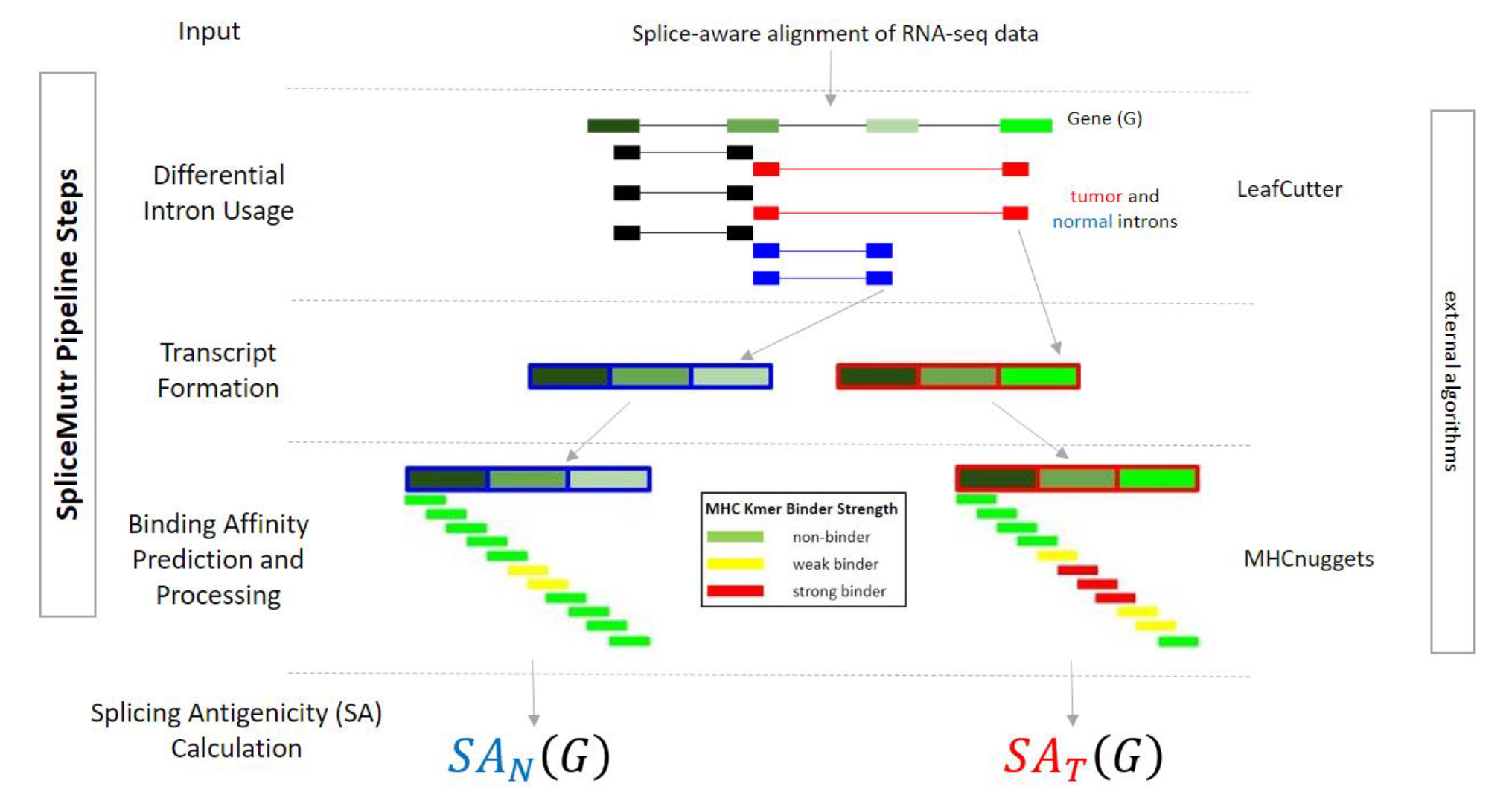
The SpliceMutr pipeline. The SpliceMutr pipeline uses RNA-seq data from two groups of samples and evaluates the changes in splicing antigenicity between them. In this example, the alternative splicing analysis compares tumor and normal RNA-seq samples. The pipeline performs splicing-aware alignment, gene expression quantification, and HLA genotyping. The splicing-aware alignment intron counts are then input into LeafCutter to evaluate differential intron usage. The tumor-specific and normal-specific introns undergo transcript formation, translation, and kmerization for MHCnuggets input through SpliceMutr, then are evaluated for genotype-specific MHC binders using MHCnuggets (Shao et al., 2020). MHC binders associated with normal-specific and tumor-specific peptides are then used to calculate a per gene and sample splicing antigenicity (SA) metric dependent on the type of the sample. *SA*_*T*_ (*G*) is calculated for tumor samples and *SA*_*N*_(*G*) is calculated for normal samples.

Finally, SpliceMutr outputs splicing antigenicity metrics per gene and per sample to identify neoantigen candidates and compare splicing antigenicity with respect to other categorical or covariate data related to the genes or samples of interest. The splicing antigenicity is a measure of the potential for tumor-specific or normal-specific splicing variants to create antigens, and the tumor splicing antigenicity is normalized by the tumor purity. The higher the tumor splicing antigenicity of a gene of interest, the more likely the gene of interest harbors splicing variants with the potential for being useful antigens. We note that these metrics and, thus, downstream analysis depend on the specific algorithms selected for analysis in the pipeline. Our choice of LeafCutter (Y. I. Li et al., 2018) and MHCNuggets (Shao et al., 2020) was based on LeafCutter’s benchmarking for differential splicing and MHCNuggets’ benchmarking involving MHC-binding prediction pan-cancer, respectively. Still, we note that the pipeline can be extended to other algorithms for differential splicing and antigen prediction. Nonetheless, the use of these robust tools enables us to apply the metrics resulting from the complete SpliceMutr pipeline to evaluate the downstream impacts of splicing on the immune landscape of tumors in large-scale cohorts of short-read RNA-sequencing data in this study.

### The splicing antigenicity metric

The gene splicing antigenicity metric is calculated as follows:

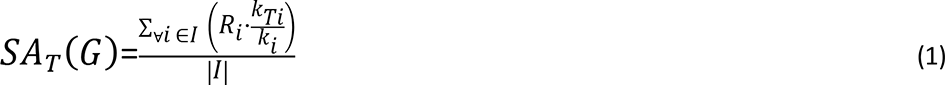

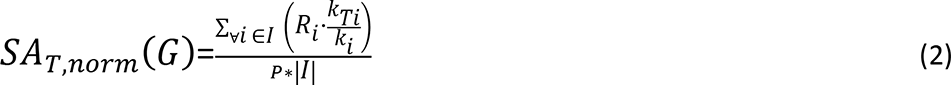

where *SA*_*T*_(*G*) is the tumor splicing antigenicity for gene *G*, *R*_*i*_ is the variance-stabilized intron read count, *k*_*Ti*_ is the number of tumor-specific binders from the peptide associated with the transcript modified by intron i, *I* is the number of tumor-specific introns for gene *G*. Using parameters obtained from normal samples and normal sample splicing patterns, a normal splicing antigenicity, *SA*_*N*_ (*G*). *SA*_*T*,*norm*_(*G*) is the tumor purity normalized tumor splicing antigenicity for gene G where P is the tumor purity. The *SA*_*N*,*norm*_(*G*) is calculated using a tumor purity of 1. After normalization, the splicing antigenicity score can then be used to calculate a differential splicing antigenicity per gene between normal and tumor splicing patterns using a log ratio, it can be used to create a splicing antigenicity gene rank per sample, and it can be averaged to create a splicing antigenicity score per sample across all genes. From here on out the splicing antigenicity will refer to equation (2).

## Results

### Splicing antigenicity displays widespread pan-cancer variability and is anti-correlated with tumor mutational burden

Defining splicing antigenicity as a sample-specific metric through SpliceMutr enables the comprehensive evaluation of its impact on the tumor immune landscape through pan-cancer analysis in the Cancer Genome Atlas (TCGA). To evaluate the prevalence of splicing antigenicity in a pan-cancer manner, we first evaluated the tumor vs the normal splicing antigenicity per cancer subtype (Fig. 2A). Among the tumor types analyzed, we observed a trend towards increased splicing antigenicity in tumor vs normal samples pan-cancer, which is statistically significant (measured by Wilcoxon) and higher in tumor samples for BLCA (p-value=1.10E-4, Cohens d=-0.28), BRCA (p-value=5.80E-12, Cohens d=-0.17), COAD (p-value=1.41E-2, Cohens d=0.06), HNSC (p-value=2.6E-7, Cohens d=-0.15), KICH (p-value=2.8E-4, Cohens d=-0.71), KIRC (p-value=1.39E-2, Cohens d=0.15), LIHC (p-value=4.52E-2, Cohens d=0.01), LUAD (p-value=2.5E- 8, Cohens d=-0.07), LUSC (p-value=5E-5, Cohens d=0.06), PRAD (p-value=2.5E-16, Cohens d=- 0.55), READ (p-value=1.47E-2, Cohens d=-0.27), THCA (p-value=8.8E-16, Cohens d=-0.71), and UCEC (p-value=1.2E-15, Cohens d=-0.65). Many cancers have negligible effect sizes (COAD, LIHC, LUAD, LUSC), but all tumor subtypes with significant differences in splicing antigenicity across tumor and normal samples have a higher splicing antigenicity in the tumor samples.

**Fig. 2.**
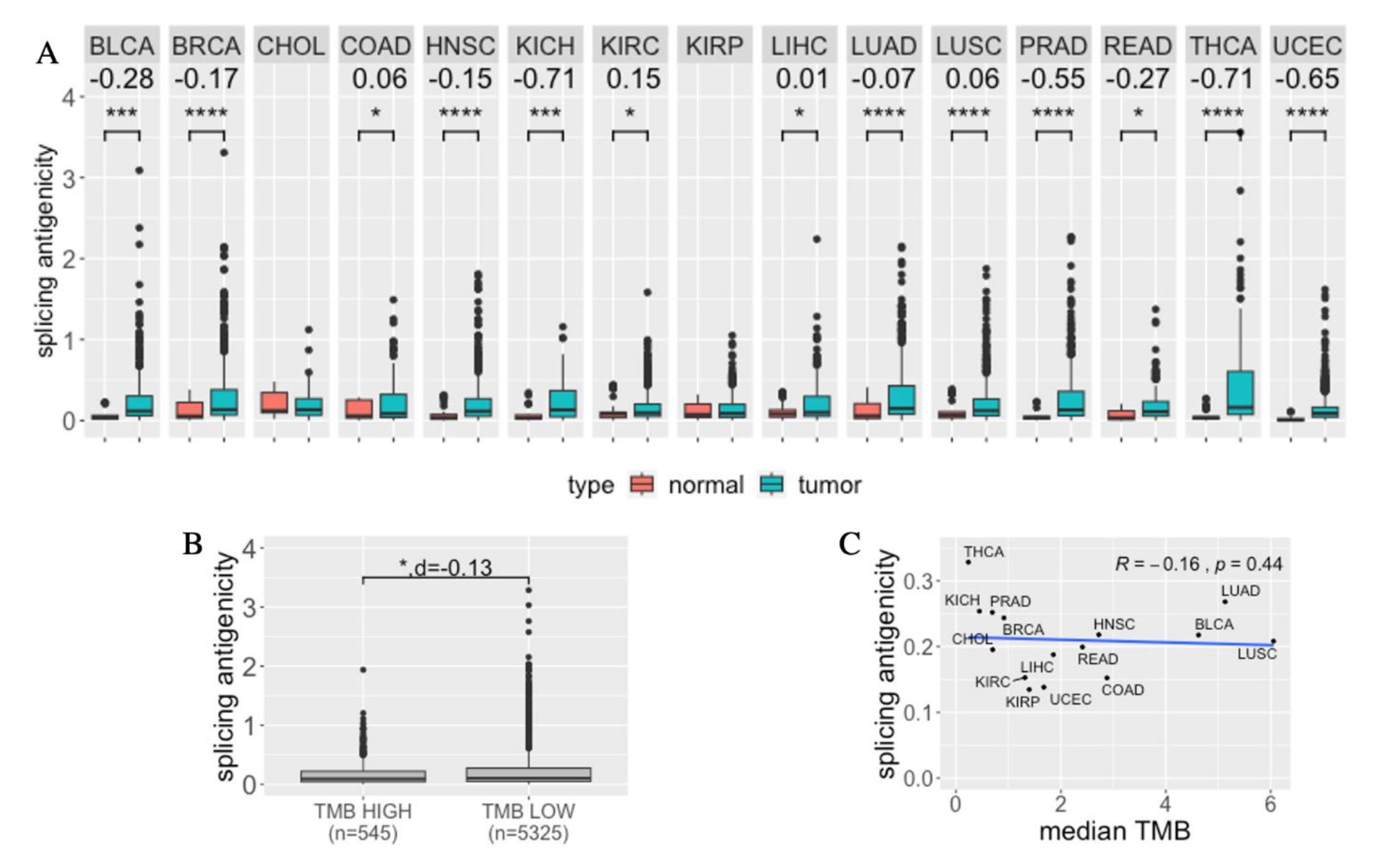
Splicing antigenicity by tissue type and in relation to the tumor mutational burden. (A) The splicing antigenicity averaged across genes per sample for tumor and normal samples for each tumor type analyzed (Wilcoxon test, Cohens d). (B) The splicing antigenicity averaged across all genes per sample for TMB high and TMB low samples for all analyzed TCGA samples (Wilcoxon test, Cohens d). (C) The per TCGA tumor type plotted against the splicing antigenicity averaged across genes per sample, then sample per TCGA cohort (Kendall Tau test). * p-value<=0.05, ** p-value<=0.005, *** p-value<=0.0005, **** p-value<=0.00005. See TCGA cancer type abbreviations in Table S1.

The observation of increased tumor splicing antigenicity in many cancer subtypes leads us to hypothesize that the total level of splicing antigens in a tumor may impact its immunogenicity and immunotherapy response. To evaluate the connection of splicing antigenicity to established biomarkers for immunotherapy pan-cancer, we next stratified all TCGA samples in TMB high (TMB >= 10 mutations/Mb) and TMB low (TMB < 10 mutations/Mb) subgroups (Riviere et al., 2020; Wu et al., 2022) and quantified differences in their splicing antigenicity averaged across genes per sample (Fig. 2B). We find that TMB high tumors have a significantly lower splicing antigenicity than TMB low tumors, although the Cohens d is small (p-value = 0.024, Cohens d=-0.13, Wilcoxon test). To further test this hypothesis, we explored the association of splicing antigenicity per tumor type across all samples to TMB (Fig. 2C). In this comparison, we found that the median splicing antigenicity per tumor type in TCGA is insignificantly negatively correlated to the median TMB per tumor type in TCGA (p-value=0.44, tau=-0.16, Kendall Tau Test). Altogether, these findings support the conclusion that tumor-associated splicing variants have more immunogenic potential than normal, and that the splicing antigenicity is independent of the TMB. These results warrant further analysis within tumor subtypes to determine the impact of tumor splice variants on immunogenicity.

### The abundance of splicing antigens associates with the frequency of mutations in the splicing machinery

Mutations in the catalytic RNA-core (Inoue et al., 2019; Zhang et al., 2019) and in scaffold splicing factor proteins (Kouyama et al., 2019; Liu et al., 2020b; Maguire et al., 2015; Shuai et al., 2019) have been associated with splicing alterations. To examine potential links between mutations in the splicing machinery with splicing antigenicity across tumor types, we first categorized samples in TCGA based on the presence of non-silent mutations in splicing factor genes SF3B1 (Shirai et al., 2017) and analyzed levels of splicing antigenicity (Table S2). The only cancer subtypes that showed significant differences in splicing antigenicity were between SF3B1-wt and mutant samples in BRCA and HNSC (Fig 4A-4B, Table S2), with higher splicing antigenicity in mutant samples (BRCA p-value = 0.047 and Cohens d = 0.4; HNSC p-value =0.0021 and Cohens d = 0.83, Wilcoxon test). This observation is consistent with the hypothesized role of SF3B1 as a biomarker for immunotherapy currently under evaluation in a breast cancer clinical trial (ClinicalTrials.gov, NCT04447651) but suggests that it may not generalize to additional cancer subtypes except for head and neck cancer.

**Fig. 4.**
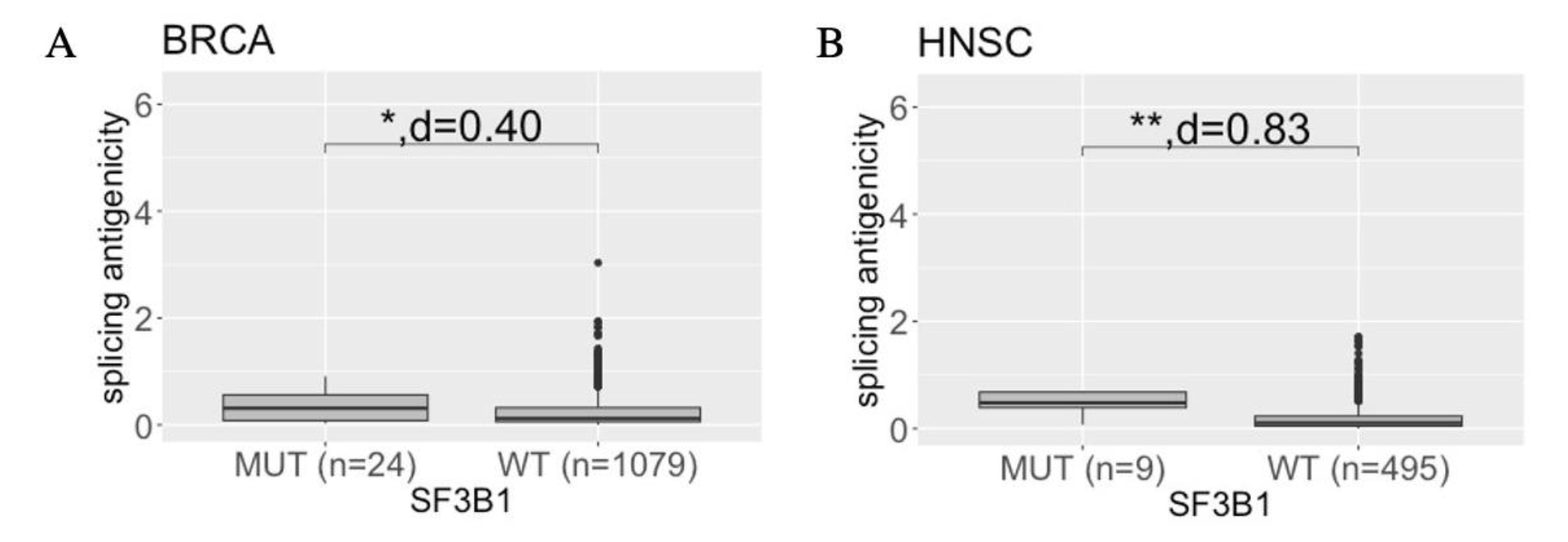
Pan-cancer correlation analysis of per-sample splicing antigenicity with non-silent mutations in splicing machinery coding genes. Distribution of splicing antigenicity per sample, averaged across genes, by mutations in the splicing factor gene SF3B1 in BRCA (A) or HNSC (B). * p-value<=0.05, ** p-value<=0.005, *** p-value<=0.0005, **** p-value<=0.00005. See TCGA cancer type abbreviations in Table S1.

### Response to immunotherapy in melanoma patients is associated with decreased splicing antigenicity

While the TCGA analysis enables us to correlate splicing antigenicity to various biomarkers of ICI response pan-cancer, including TMB, this database is only composed of pre-treatment samples and does not allow the direct comparison of splicing antigenicity before and after treatment. Therefore, to directly associate splicing immunogenicity with ICI response, we applied SpliceMutr to a cohort of RNA-seq data from 67 metastatic or advanced melanoma patients treated as part of an ICI clinical trial described previously (ClinicalTrials.gov, NCT01621490, Grasso et al. 2020; Anagnostou et al. 2020). Briefly, this cohort has RNA-seq data pre-and post-treatment in patients from three arms. Patients in the NIV1+IPI3 arm were treated with Nivolumab (anti-PD1) and Ipilimumab (anti-CTLA4) in combination (n=8 patients) and had no previous exposure to Ipilimumab. In the NIV3-PROG arm, patients were previously exposed to Ipilimumab and were only treated with Nivolumab after progression while on Ipilimumab (n=32 patients). In the NIV3-NAIVE arm, patients had no previous anti-CTLA4 therapy and were only given Nivolumab as a monotherapy (n=27 patients).

This melanoma cohort contains only tumor samples, but not their non-cancer controls, limiting an analogous comparison of differential splicing used in the TCGA. To account for the smaller sample size and a lack of normal controls in this melanoma cohort, we used LeafCutterMD (Jenkinson et al., 2020) to perform a one vs ipilimumab-naive baseline outlier analysis (Fig. S3, Fig. S4). The outlier analysis method using LeafCutter makes comparing splicing differences between conditions with small sample sizes, in contrast to the requirement of 6 replicates per condition in the differential splicing analyses used for TCGA. This outlier splicing analysis is useful for early-phase clinical trial analysis where there are typically a small number of responders for comparison against non-responders. Using LeafCutterMD, we identified 6674 total genes impacted by outlier splicing. Using this subset of genes, we calculated the mean splicing antigenicity per sample, and we used the Wilcoxon test p-values to determine significant differences in splicing antigenicity within treatment arm and across objective response (Fig. 5). The CRPR group are all complete and partial responders, the SD group are all stable disease patients, and the PD group are all progressive disease patients.

**Fig. 5.**
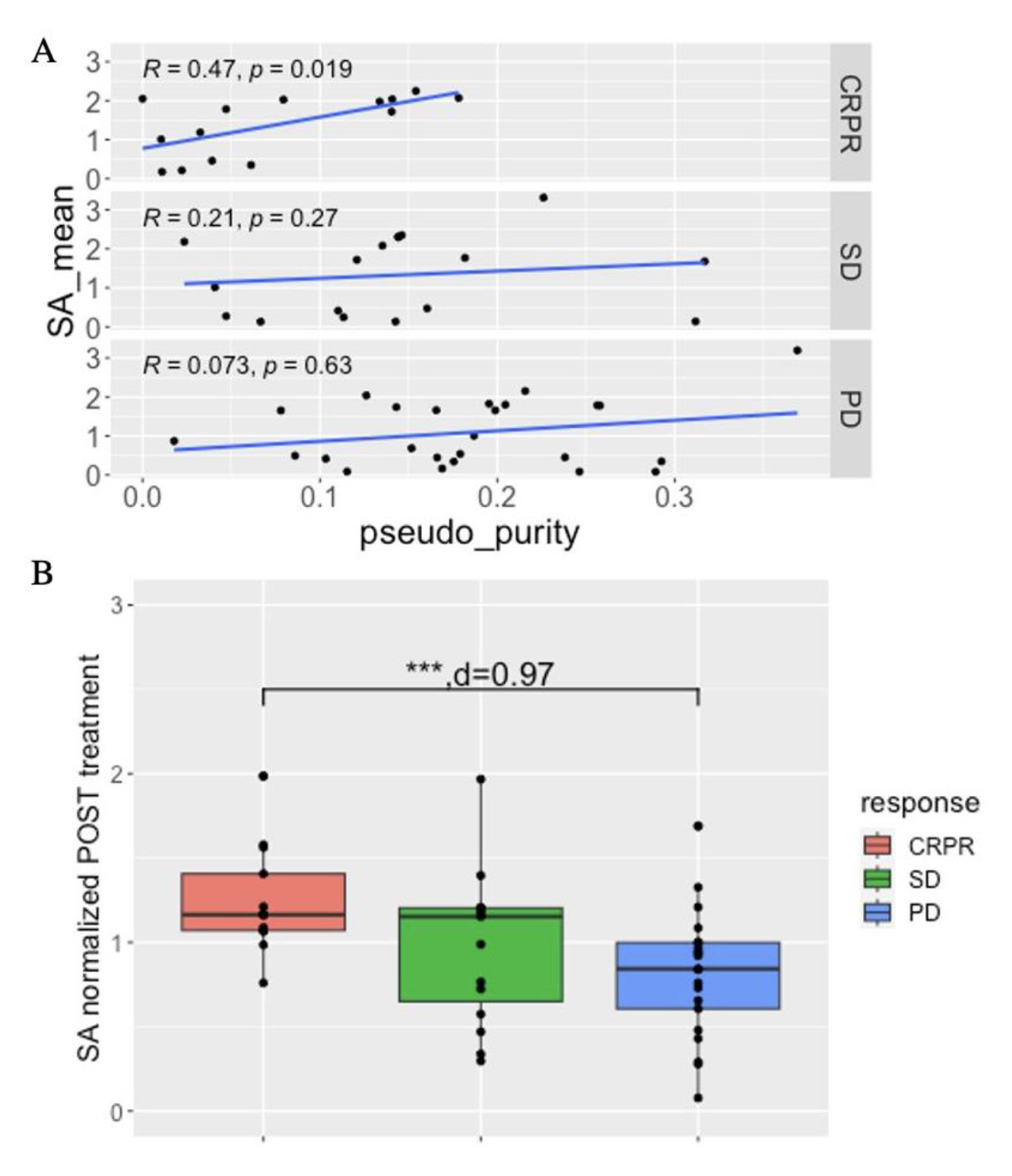
Per-patient splicing antigenicity of ICI-treated melanoma patients per treatment arm and response type after treatment. (A) The per-patient pseudo purity compared to the mean splicing antigenicity averaged across genes by the response for all treatment arms. Kendall Tau test. (B) The mean splicing antigenicity averaged across genes per patient and normalized by the pseudo purity, for all treatment arms combined. Wilcoxon test and Cohens d. * p-value<=0.05, ** p-value<=0.005, *** p-value<=0.0005, **** p-value<=0.00005.

Whereas TCGA tumors were obtained from surgical biospecimens, and quality controlled to ensure tumor purity prior to sequencing, this clinical trial uses biopsies for sequencing. The resulting variability in tumor purity in these biospecimens may impact the splice variants detected and subsequent estimates of splicing antigenicity. To evaluate the impact of the therapeutic effect on the splicing antigenicity metric, we plotted the ESTIMATE-calculated pseudo purity (Yoshihara et al., 2013) vs the splicing antigenicity (Fig 5A). We found that the pseudo purity was significantly positively correlated with the splicing antigenicity in responders (Kendal Tau test, p-value=0.019, tau=0.47). Therefore, we decided to normalize the splicing antigenicity, per sample, by the pseudo purity by modifying equation (1) to equation (2). Once normalized, we evaluated the splicing antigenicity independent of treatment arm across response (Fig. 5C). We found that responders had a significantly higher splicing antigenicity than progressive disease patients (Wilcoxon test, p-value= 1.2E-4, Cohens d=0.97; Fig. 5C). Although we do not see the same trends prior to treatment (Fig. S2), significance in responders, at least for the NIV-PROG arm is not reached due to outlier samples and a minimal number of responders available for comparison. These results show that, once normalization is done for therapeutic effect, responders have a higher immunogenic potential.

To validate the pairwise relationship between the normalized splicing antigenicity and response to ICI therapy in this melanoma cohort, we extracted the splice junctions associated with the union of the top twenty genes, by splicing antigenicity, impacted by outlier splicing per sample. Then we averaged the splicing antigenicity per splice junction across samples by response to obtain a mean splicing antigenicity per splice junction. We then performed a pairwise Wilcoxon test between the splicing antigenicity of the same splice junctions in responders compared to baseline, stable disease patients compared to baseline, and progressive disease patients compared to baseline. We found that responders (p-value= 1.77E-294,, Cohens d = −0.85; Fig. 6, Fig. S6) and stable disease patients (p-value= 9.45E-247, Cohens d = −0.29; Fig. 6, Fig. S6) have a significantly increased splicing antigenicity compared to baseline while progressive disease patients (p-value= 5.33E-47, Cohens d = 0.14; Fig. 6, Fig. S6) have a significantly decreased splicing antigenicity compared to baseline. Further, there is a decreasing splicing antigenicity as response worsens (Fig. 6, Fig. S6). These results further suggest that the response in this cohort is in part mediated by the immunogenic potential of splicing antigens.

**Fig. 6.**
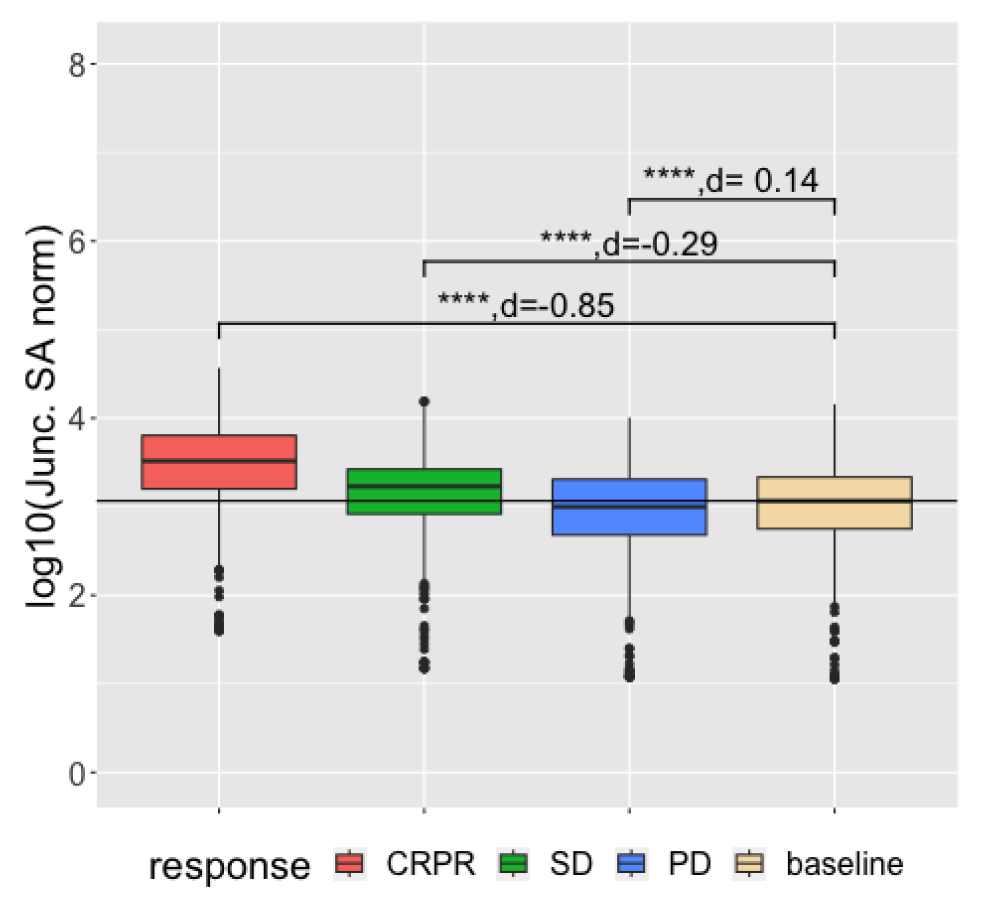
Per splice junction normalized splicing antigenicity for responders compared to baseline samples. The splicing antigenicity averaged across samples for the subset of splice junctions derived from the top twenty genes with the highest splicing antigenicity per sample. The black horizontal line is the median splicing antigenicity for baseline samples. * p-value<=0.05, ** p-value<=0.005, *** p-value<=0.0005, **** p-value<=0.00005.

## Discussion

This pan-cancer evaluation of splicing antigenicity using SpliceMutr demonstrates that the splicing antigenicity is higher in tumor samples than in normal samples and is weakly and insignificantly negatively correlated with TMB. Additionally, this study shows that in some tumor types, mutations in the splicing machinery significantly alter the tumor splicing antigenicity of samples that harbor these mutations. The evaluation of SpliceMutr splicing antigenicity in the melanoma cohort treated with immune checkpoint inhibitor therapy shows that the tumor splicing antigenicity is increased in responders relative to non-responders and that responders are characterized by increased splicing antigenicity per splice-junction after ICI therapy relative to before.

Our evaluation of splicing antigenicity using the RNA-seq data from immunotherapy-treated melanoma samples in the CM-038 clinical trial (ClinicalTrials.gov, NCT01621490, Grasso et al. 2020; Anagnostou et al. 2020) demonstrates that splicing antigenicity is significantly increased in responders relative to non-responders. This observation is inconsistent with the weak significant anti-correlation of splicing antigenicity to TMB, although this significance is driven by the outlier TCGA cancer subtype thyroid carcinoma. When removing thyroid carcinoma, there is a weak and insignificant positive correlation between the splicing antigenicity and TMB. Further, our work shows that those patients that respond well to treatment have a larger difference between the splicing antigenicity after treatment relative to baseline compared to those that respond poorly. These findings are contradictory to the findings of Eyras’s lab (Trincado et al., 2021), where their calculation of the number of splicing-derived tumor neoantigens showed no significant variation in the response in a cohort of melanoma patients treated with ICI therapy. It is noteworthy that although SpliceMutr uses the number of splicing-derived neoantigens as a basis for the splicing antigenicity score, SpliceMutr generates a sample-level splicing antigenicity score dependent on several additional and relevant aspects of the splicing neoantigens other than just the number. This may contribute to the discrepancy between Eyras’s group and this work. Additionally, in the Lu et al paper (Lu et al., 2021), they find that modulation of splicing factor function in combination with ICI therapy results in the development of splicing-derived neoantigens that can cause an immune response. Our finding that mutations in the splicing factor machinery can result in increased splicing antigenicity in some cancers relative to wild-type samples is complemented by this study. As a result, our study shows that splicing antigenicity can be increased in some cancers even without drugged modulation of the splicing factor machinery. Altogether, these findings point to a model in which splicing contributes to the antigen landscape in tumors, thereby increasing the ability of immune recognition and immunotherapy response.

Evaluating splicing neoantigens in cancer has been explored by several key studies (Kahles et al., 2018; Lu et al., 2021; Shirai et al., 2017; Trincado et al., 2021). Yet, each method focuses solely on the number of splicing-derived neoantigens produced and uses an outlier-based approach dependent on tissue-matched GTEx (Aguet et al., 2020) samples as reference (Kahles et al., 2018), an outlier-based approach dependent on the entire set of GTEx samples as normal (Trincado et al., 2021), or an approach based on the differential isoform usage of reference isoforms (Lu et al., 2021). The analysis of an independent reference in these previous studies may introduce batch effects in the evaluation of splicing antigens. Still, further work is needed to make larger cohorts of normal samples generated in the same technical batch and with true normal samples that may not be subject to the splicing alterations resulting from field effects in matched normal samples.

The SpliceMutr pipeline benefits from the introduction of several new elements to the analysis of splicing-derived neoantigens. SpliceMutr uses LeafCutter differential intron analysis as well as LeafCutterMD outlier splicing analysis to identify dysregulated splicing targets in cancer. By using these tools, SpliceMutr leverages the strengths of short-read RNA-seq: primarily high coverage, minimizes the pitfalls of short-read RNA-seq: non-multi-exon-spanning reads, and allows for the response of samples across conditions. SpliceMutr also introduces calculations of nonsense-mediated decay and subsequent filtering of non-coding transcripts into the evaluation of the alternatively spliced or outlier spliced junctions. Finally, SpliceMutr enables calculation of a tumor-purity-normalized splicing antigenicity score that enables evaluation of the impact of a tumor’s immunogenicity and can be used as an immunotherapy biomarker.

Though useful for evaluating the impact of splicing on the neoantigen burden, this study does have limitations. The SpliceMutr pipeline does not perform transcriptome assembly, instead transcripts are modified by an individual splice-junction of interest. This does not allow for multiple alternative or non-canonical events to be included in a single transcript. Additionally, SpliceMutr does not include analysis of the intron-retention alternative splicing event in differential analysis or transcript formation. This is due to the desire to perform differential and outlier intron usage analysis using LeafCutter and LeafCutterMD, each of which defines splice-junctions as exon-spanning reads. Although, it might be possible to identify retained introns using tools such as IRFinder (Middleton et al., 2017) and iREAD (H.-D. Li et al., 2020), label retained introns using codes, and then combine the identified intron-retention splice-junctions with exon-spanning splice-junctions as input into LeafCutter and LeafCutterMD. This would result in differential or outlier splicing analysis that includes intron retention while using the LeafCutter or LeafCutterMD framework. Future work extending this approach to splicing inferred from long-read sequencing may overcome some of the limitations of this pipeline resulting from the reliance on inferred transcripts from short read sequencing. While developing this pipeline for bulk RNA-seq data enables analysis of large-scale tumor cohorts, low tumor purity or mixtures of additional cell types can impact our estimates of splice variants. In this study, we address this by scaling our estimate of splicing antigenicity by tumor purity. Single-cell profiling can overcome this need for scaling by isolating tumor cells for transcriptional profiling but estimates of splice variants from this technology remain challenging due to high dropout of RNA signal. Therefore, future studies relying on profiling of malignant epithelial cells as through laser capture microdissection or improved high-coverage single cell technologies are essential to overcome these limitations. A further limitation of this study is that SpliceMutr relies on universal MHC binding affinity thresholds to establish antigens. Using MHC-specific IC50 thresholds when predicting peptide binders improves the accuracy of the prediction (Paul et al., 2013). Yet, due to using a uniform threshold for peptide binders, less than or equal to a 500 nM IC50 score, SpliceMutr does not allow for custom MHC-specific binder classification. Custom MHC-specific binder classification can be implemented by including an internal lookup table to appropriate MHC-specific binder thresholds during the binding affinity prediction and selection phase. Additionally, when making MHC:peptide predictions, MHCnuggets (Shao et al., 2020) uses either the input allele’s prediction model if it exists, or the allele closest to the allele input if the model does not exist.

## Conclusions

SpliceMutr shows that splicing antigenicity changes in response to ICI therapies and that native modulation of the splicing machinery through mutations increases the contribution of splicing to the neoantigen load of some TCGA cancer subtypes. Future studies of the relationship between splicing antigenicity and immune checkpoint inhibitor response pan-cancer are essential to establish the interplay between antigen heterogeneity and immunotherapy regimen on patient response.

## Methods

### Preprocessing RNA-seq data

SpliceMutr requires splicing-aware aligned RNA-seq data as an input. In this study, all data was aligned using STAR version 2.7.3a using the basic two pass mode and six threads (Dobin et al., 2013). Both the .bam and the SJ.out.tab files are used moving forward in the analysis. Each .bam file is used for sample genotyping by arcasHLA (Orenbuch et al., 2020). The SJ.out.tab file is a file containing a set of high-confidence splice junctions output by the STAR aligner. This SJ.out.tab file is first filtered to contain only canonical splice motifs, then the data from the SJ.out.tab file is transferred to a .junc file format that is readable by LeafCutter (Y. I. Li et al., 2018) and LeafCutterMD (Jenkinson et al., 2020) using custom scripts available from https://github.com/FertigLab/splicemute.git.

### Running LeafCutter

Each .junc file, as well as the metadata necessary to run LeafCutter (Y. I. Li et al., 2018) differential junction usage analysis, are created using custom scripts and by hand. Initially, LeafCutter creates a set of clusters of splice junctions that overlap based on their coordinates and the existence of at least three reads, providing evidence of the existence of the splice junction in the data. The script that performs this analysis is modified to accept the non-BED formatted .junc files created in the preprocessing stage. After clustering this way, we perform differential splice junction usage analysis using LeafCutter with at least six samples per comparison.

### Running LeafCutterMD

LeafCutterMD (Jenkinson et al., 2020) is an algorithm based on LeafCutter that evaluates outlier usage of splice junctions in a one-vs-many manner. Splice junction clustering is done exactly the same as it is when running LeafCutter differential splice-junction usage analysis. The difference when running LeafCutterMD is the script that processes the output of this clustering result as well as the metadata generated to perform the analysis. The outlier analysis is done in a one-vs-many fashion, so the files that determine sample groupings contain a single sample belonging to the target group and then a larger number of samples belonging to the comparator group. The splice junctions identified as having outlier expression are those of the target group having outlier usage when compared to the appropriate splice junction cluster of the comparator group. Creating the many target and comparator groups to be run by LeafCutterMD as well as collapsing the many one-vs-many outlier analysis done by LeafCutter, is carried out using custom analysis scripts.

### Sample Genotyping arcasHLA

Sample genotyping using arcasHLA is carried out by first running the extract functionality of arcasHLA on the STAR alignment output BAM files and then running the genotype functionality of arcasHLA on the output from the extract. The extract functionality of the arcasHLA toolset extracts reads that map to chromosome 6 of the human genome. The genotype functionality of arcasHLA gives the genotype of the sample given the reads mapped to chromosome 6 of the alignment file. The genotype per sample is then extracted and saved for per sample MHC:peptide binding affinity predictions while running that portion of SpliceMutr.

### Running MHCnuggets

The per-sample genotype output from arcasHLA as well as the tumor-specific and normal-specific kmers output from SpliceMutr, are input into MHCnuggets (Shao et al., 2020). Each kmer undergoes binding affinity prediction using each MHC-class 1 allele from each sample-specific genotype to determine a set of tumor-specific and normal-specific antigenic kmers.

### The LGC (ORF Length and GC content) Coding Potential Calculation

Briefly, the modified LGC coding potential (Wang et al., 2019) is calculated as follows:

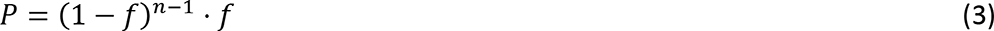

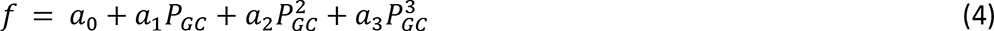

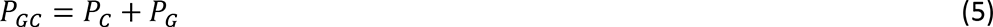

where *P*_*C*_ is the probability of cytosine, *P*_*G*_ is the probability of guanine, *P* is the probability of the ORF in the sequence, *f* is the probability of finding a stop codon in the sequence, *n* is the number of sense codons in the sequence, and the parameters *a*_0_through *a*_3_are coefficients learned from protein-coding human transcripts or long non-coding human transcripts. This coding potential can then be used to calculate the differential coding potential (L) between the coding LGC score (P_c_) and the long non-coding LGC score (P_nc_) for the same transcript:

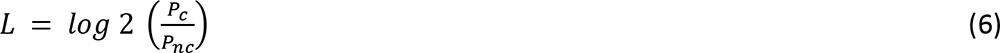

### Calculating the pseudo purity

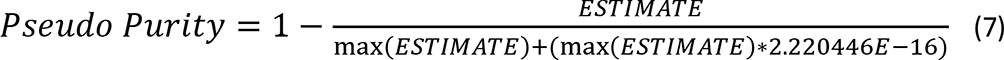

The pseudo purity is calculated for a cohort using the ESTIMATE (Yoshihara et al., 2013) score normalized by the maximum ESTIMATE score for the cohort plus the product of the maximum and machine epsilon in the R statistical package. This is so that the maximum ESTIMATE score for the cohort does not result in a sample with a tumor purity of zero. Only samples with tumor purity greater than 0.01 are used for analysis.

### Calculating the differential agretopicity

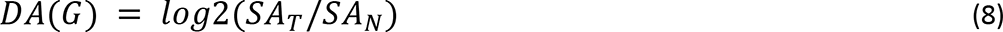

The differential agretopicity (Richman et al., 2019) per gene is calculated as the log2 ratio of the tumor splicing antigenicity per gene divided by the normal splicing antigenicity per gene.

### The SpliceMutr Pipeline - Intron Parsing

At its basic level, the SpliceMutr toolset takes as input a set of introns in BED file format, their chromosome location, and splice site coordinates. Per intron, SpliceMutr identifies whether the intron has a pair of splice sites that are found spliced together in the associated reference transcriptome. If so, the intron is classified as an annotated intron and is classified as an unannotated intron if not. If the set of introns in the BED file has been previously annotated, as is the case with LeafCutter analysis, SpliceMutr will carry those specific annotations through to further analysis. LeafCutter-specific annotations of splice sites are detailed in Supplemental Fig. S1. Given supplied gene annotations, LeafCutter will identify specific splice-junctions with respect to the reference. LeafCutter defines an annotated splice-junction as a splice-junction with both 5’ and 3’ splice sites matching known reference splice sites that are annotated as being spliced together. A cryptic splice-junction is defined as having a 3’ splice site but not a 5’ splice site, a 5’ splice site but not a 3’ splice site, or both 3’ and 5’ splice sites that do not match known reference splice sites. A novel annotated splice-junction is defined as a splice-junction with 5’ and 3’ splice sites that match known reference splice sites that are not annotated as being spliced together.

### The SpliceMutr Pipeline - Transcript Identification and splice-site modification

After intron-parsing, the next step in the SpliceMutr pipeline is to identify the pair of flanking exons associated with each splice site for the target splice junction. Using each individual flanking exon, the upstream flanking transcripts are selected if they contain the upstream flanking exon, and the downstream flanking transcripts are selected if they contain the downstream flanking exons. All combinations of a single upstream transcript and a single downstream transcript are formed. In the case of an intron that is classified as annotated, only flanking transcripts with matching annotation names are preserved within the transcript combinations. Additionally, since the intron is annotated, the flanking exons should be directly spliced together within the transcript. If this is not the case, the matching flanking transcript pair is not preserved. With the unannotated introns, all transcript pairings are preserved unless it is determined that there is a subset of matching transcript pairings that contain directly adjacent flanking exons. In this case, only those matching transcript pairings with directly adjacent flanking exons are preserved for analysis.

Once a set of candidate flanking transcript pairings are identified, transcript modification begins. To perform transcript modification, the full spliced upstream flanking transcript is joined to the full spliced downstream flanking transcript using the splice site coordinates of the target splice-junction as the joining point. If both the upstream and downstream flanking transcript have annotated UTRs (untranslated regions) based on the reference transcriptome, then the joined transcript product is annotated as protein-coding. This creates a joined transcript containing both five-prime and three-prime UTRs. If both the upstream and downstream flanking transcripts do not have annotated UTRs regions, then the joined transcript product is annotated as non-protein-coding. Now SpliceMutr performs open reading frame (ORF) finding and translation of each joined transcript identified as protein-coding.

### The SpliceMutr Pipeline - Open-reading Frame Finding and Coding Potential Calculation

To perform ORF finding, SpliceMutr searches for the first start codon beginning at the five-prime UTR of the joined transcript. Once the first start codon is identified, SpliceMutr searches the transcript until the first stop codon is identified. The region from the first identified start codon to the first identified stop codon is extracted as the open-reading frame for the transcript. Once the ORF is identified, the transcript is translated and saved for downstream analysis. ORF finding is performed this way to ensure that the experimentally validated open reading frame is conserved during transcript modification. All modifications to the open reading frame per target intron and joined transcript are recorded in the SpliceMutr metadata (Fig S2).

In the final stage of transcript modification, the coding potential of each target-intron-modified transcript of size greater than or equal to 100 nucleotides is calculated using a modified version of the ORF **L**ength and **GC** content (LGC) method (Wang et al., 2019). Only those transcripts with a coding potential greater than zero are used in subsequent analysis.

### The SpliceMutr Pipeline - MHC Binding Affinity Predictions and alternative binder filtering

The set of proteins output from transcript modification are then processed and MHC binding affinity predictions are performed on tumor-specific and normal-specific kmers. The translated proteins are kmerized, then SpliceMutr extracts out the unique set of kmers found in the set of translated proteins. SpliceMutr then uses MHCnuggets (version 2.3.2) (Shao et al., 2020) to predict the raw binding affinity of each kmer using the HLA alleles specific to the samples being analyzed. SpliceMutr then extracts out those kmers with a predicted IC50 less than or equal to 500 nM per HLA allele. Briefly, IC50 predictions are approximations of in-vitro measurements of the concentration of a target peptide necessary to inhibit the binding of a high-affinity radiolabeled peptide to a specific MHC molecule by 50 percent (Sidney et al., 2013). The lower this IC50 concentration, the higher affinity the target peptide has for the specific MHC molecule. If differential splice junction usage is calculated, predicted binders produced by tumor-specific transcripts are further filtered such that all binders also found within alternative normal-specific transcripts are removed. Predicted binders produced by normal-specific transcripts are filtered similarly to how tumor-specific binders are filtered, except all binders from the tumor-specific alternative event are filtered out. This filtering is not carried out when performing outlier splicing analysis.

### Gene and sample summaries of splicing antigenicity from RNA-seq analysis with SpliceMutr

In summary, SpliceMutr inputs sample HLA type, differentially used or outlier splice junctions, and variance-stabilized splice-junction counts to calculate the number of filtered immunogenic kmers per splice-junction-modified transcript and per sample. SpliceMutr then uses this information to calculate the per-gene splicing antigenicity metric. If differential splice junction usage is carried out, the differential agretopicity can be calculated per gene and used to filter for those genes with higher splicing antigenicity in tumor transcripts than normal. The per gene splicing antigenicity metric can be calculated, which can be further summarized per sample by taking the average across all differential or outlier splicing-impacted genes. This provides a summary of splicing antigenicity per gene and further per sample for further statistical analysis.

### TCGA Pan-Cancer Analysis

TCGA cancer subtypes were selected for analysis based on whether the cohort had at least six tumor and six normal samples with RNA-seq data. A total of 15 tumor subtypes fit this evaluation criterion: prostate adenocarcinoma (PRAD), thyroid carcinoma (THCA), kidney renal clear cell carcinoma (KIRC), lung squamous cell carcinoma (LUSC), lung adenocarcinoma (LUAD), breast invasive carcinoma (BRCA), liver hepatocellular carcinoma (LIHC), bladder urothelial carcinoma (BLCA), kidney renal papillary cell carcinoma (KIRP), colon adenocarcinoma (COAD), head and neck squamous cell carcinoma (HNSC), uterine corpus endometrial carcinoma (UCEC), rectum adenocarcinoma (READ), cholangiocarcinoma (CHOL), and kidney chromophobe (KICH). Splice-junction genomic coordinates and counts per tumor and normal sample in each cohort were extracted from recount3 (version 1.6.0) (Wilks et al., 2021) for LeafCutter (version 0.2.9) analysis (Y. I. Li et al., 2018). Those differential splice-junctions were then run through SpliceMutr to generate a set of genes and their tumor-specific splicing antigenicity scores. Optitype HLA calls were used to obtain HLA genotypes per TCGA sample (Szolek et al., 2014; Thorsson et al., 2018) for SpliceMutr analysis.

We used the multi-omic data from TCGA to further associate splicing antigenicity with common immunotherapy biomarkers. We specifically correlated the splicing antigenicity to a series of immune metrics, mutation-based metrics, and somatic mutation calls for each TCGA sample (Thorsson et al., 2018). The metrics used for correlative analysis were the TCR clonality, leukocyte fraction, number of immunogenic mutations, CIBERSORT cell type estimates (Chen et al., 2018), and the tumor mutational burden (TMB) calculated as the number of non-silent mutations per megabase in coding regions. The somatic mutation calls were used to determine mutations in 119 driver splicing factor genes per TCGA cancer subtype and sample (Seiler et al., 2018b). Using this data, we stratified TCGA samples per cancer subtype based on the existence of any non-silent mutation first in each individual driver splicing factor gene, then based on the existence of any non-silent mutation in at least one of the 119 driver splicing factor genes, and finally calculated whether the splicing antigenicity differed significantly between samples with mutations and samples without mutations using the Wilcoxon statistic and significance determined by Benjamini-Hochberg (BH) adjusted p-values below 0.05.

### Evaluating tumor-specific splicing antigenicity in a melanoma cohort treated with Nivolumab, Ipilimumab, or their combination

We obtained RNA-seq data from the previously described CM-038 melanoma immune checkpoint trial (ClinicalTrials.gov, NCT01621490, Grasso et al. 2020; Anagnostou et al. 2020). This cohort contained biospecimens pre-and post-immunotherapy treatment in three trial arms. Patients in the first trial arm were given Nivolumab and Ipilimumab in combination (NIV1+IPI3, n=8). Patients in the NIV3-PROG arm (n=32) had previous exposure to Ipilimumab treatment and were only treated with Nivolumab. Finally, patients in the NIV3-NAIVE arm (n=27) had no previous anti-CTLA4 therapy and were only given Nivolumab therapy. Patient response to treatment was evaluated using the RECIST 1.1 criteria (Eisenhauer et al., 2009). RNA-seq data for this cohort were preprocessed by and run through SpliceMutr to generate splicing antigenicity scores per gene impacted by outlier splicing. Whereas our previous studies performed differential splicing between tumor and normal samples, no normal samples were available for this cohort. Therefore, analysis of splicing differences was carried out within treatment arms with respect to RECIST 1.1 response type and treatment time using LeafCutterMD. Responder (CRPR), stable disease (SD), and progressive disease (PD) patient splice junction data were compared to baseline splice-junction data using LeafCutterMD (Fig S3, Fig. S4). The outlier splicing model in LeafCutterMD (Jenkinson et al., 2020) was carried out to allow for uniform splicing analysis due to the small number of samples per response type in this cohort. LeafCutterMD output was then run through SpliceMutr to generate splicing antigenicity scores per gene and sample using the per-sample genotype generated from arcasHLA. The union of all splice junctions identified as being outlier splice-junction per sample were combined to create the set of splice junctions for splicing antigenicity calculations. Once calculated, the splicing antigenicity was normalized by pseudo purity, but only those samples with pseudo purity greater than 0.01 were used for analysis. This resulted in dropping the lowest purity sample from analysis. When comparing the splicing antigenicity within treatment arm and across response type, averaged across genes or samples, the Wilcoxon test statistic with BH-adjusted p values (significance <= 0.05) was used. The non-adjusted Wilcoxon test-statistic was used when comparing the splicing antigenicity across response type disregarding treatment arm. The paired Wilcoxon test was not used for this analysis because it would require missing gene-splicing antigenicity values across conditions to be filled in with zeroes. The Cohens d is calculated similar to the effect size, except it uses a pooled standard deviation (Lakens et al., 2013).

### Statistical analysis

All statistical analyses in this paper are performed in R (R Core Team). When testing for significant variation in the splicing antigenicity across conditions, the Wilcoxon test statistic with p values (significance <=0.05) was used. Multiple testing (Benjamini-Hochberg) correction was performed when greater than three comparisons are made. When testing the correlation between the splicing antigenicity and other metrics of response to immunotherapy, the Kendall Tau test statistic (significance <= 0.05) was used.

### Availability of Data and Materials

All TCGA data used for analysis in this paper were recount3 (Wilks et al., 2021) splice-junction counts and metadata and The Immune Landscape of Cancer (Thorsson et al., 2018) open and protected supplementary data.

RNA-sequencing data used for analysis of the melanoma cohort can be found at EGAS00001004545 (deposited in the European Genome phenome Archive) and human RNA sequencing data are deposited into a controlled-access database managed by Bristol-Myers Squibb. Sequencing data may be accessed, on approval, by individual investigators upon request (https://fasttrack.bms.com and).

The SpliceMutr pipeline is available from https://github.com/FertigLab/splicemute.git. All analyses in this paper can be reproduced using the scripts and directions in this repository. All figures in this manuscript can be produced from https://github.com/FertigLab/splicemutr_paper.git.

## Supplement

**Table S1.**
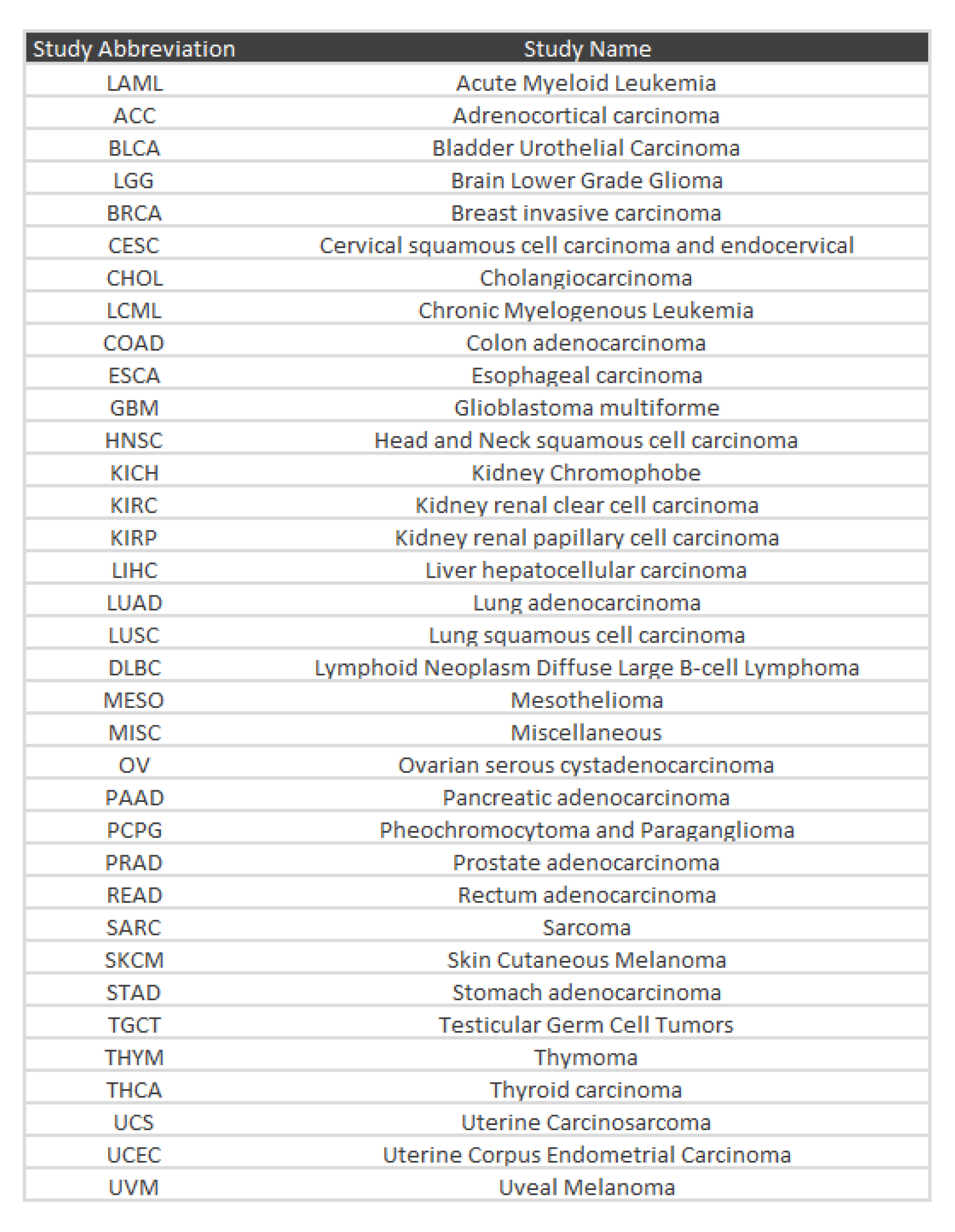
**TCGA cancer study abbreviations and names.**

**Fig S1.**
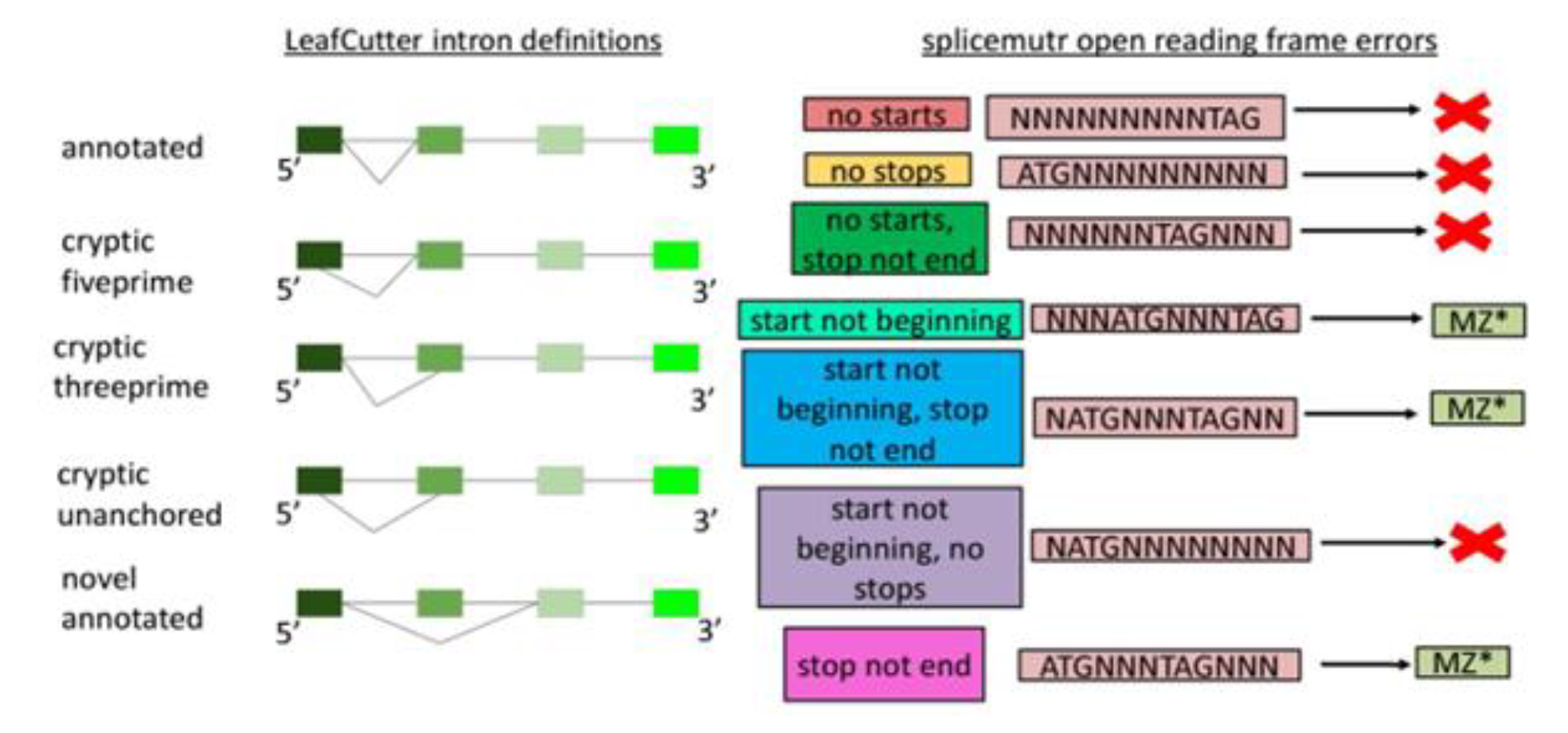
**LeafCutter intron definitions and SpliceMutr open reading frame errors.** The LeafCutter intron definitions are a visual representation of the LeafCutter intron definitions with respect to a mock gene model. Annotated introns have a pair of splice sites located at the genomic coordinates of a documented intron. Cryptic intron types have either one or both splice sites located at genomic coordinates not associated with a documented intron. Novel annotated introns have a pair of splice sites individually located at the genomic coordinates of documented introns but not at splice site locations documented to be joined together. The splicemutr open reading frame errors are a visual representation of the open reading frame (ORF) errors that splicemutr documents during transcript formation. The full range of the DNA equivalent of stop codons are defined as such during transcript formation, but this example only features the TAG stop codon. If an open reading frame can be found after modification of a reference transcript by a differentially used intron, then the transcript is translated with the appropriate error documented. If the modified transcript has unchanged ORF start and end codons relative to the reference, then the modified transcript is documented as being immediately translatable.

**Fig. S2.**
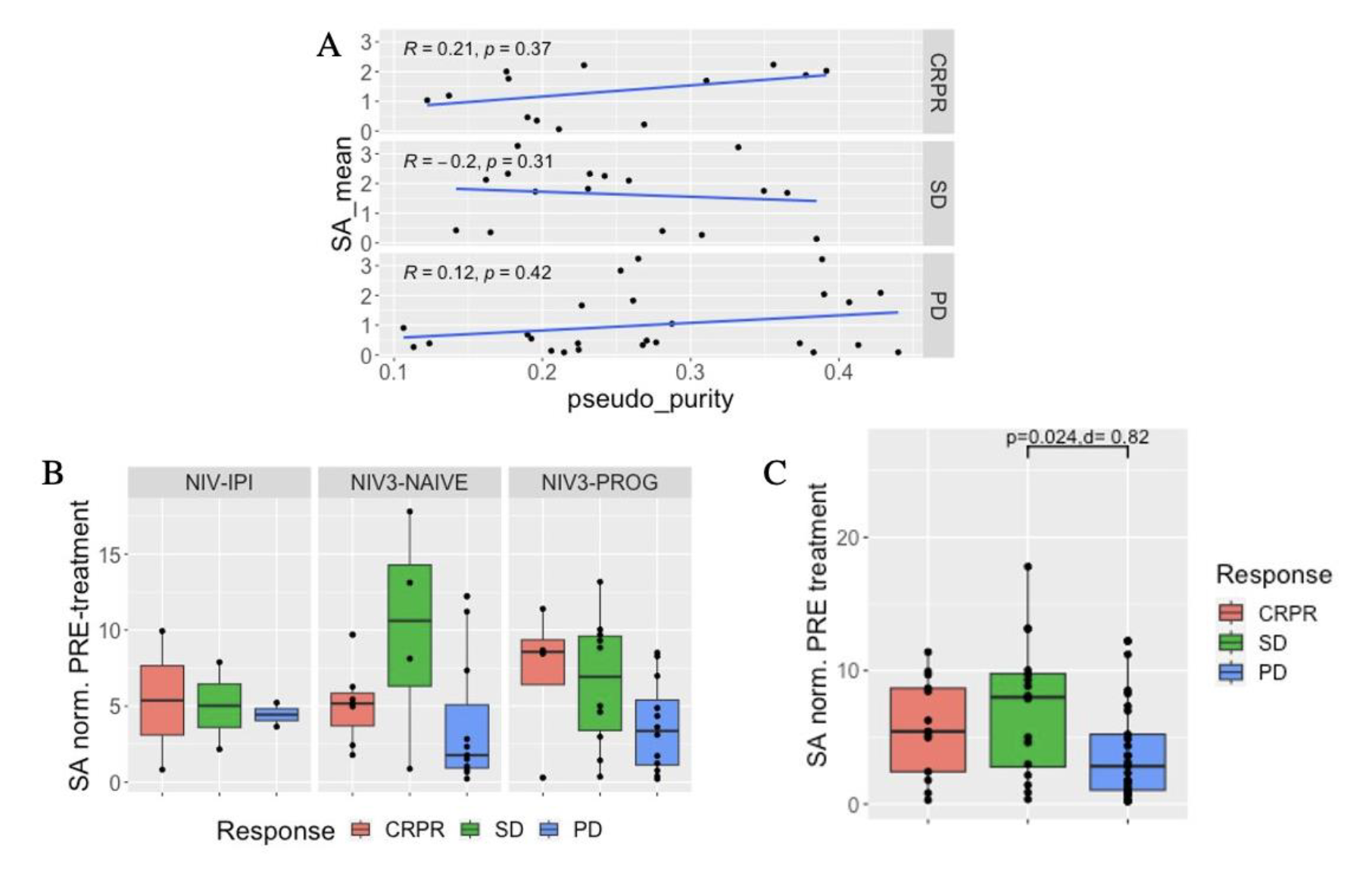
**Per-patient splicing antigenicity of ICI-treated melanoma patients per treatment arm and response type before treatment.** (A) The per-patient pseudo purity compared to the mean splicing antigenicity averaged across genes per-response type for all treatment arms. Kendall Tau test. (B) The mean splicing antigenicity averaged across genes per patient and normalized by the pseudo purity, for each treatment arm. Wilcoxon test with false discovery rate adjustment and Cohens d. (C) The mean splicing antigenicity averaged across genes per patient and normalized by the pseudo purity, for all treatment arms combined. Wilcoxon test and Cohens d.

**Fig. S3.**
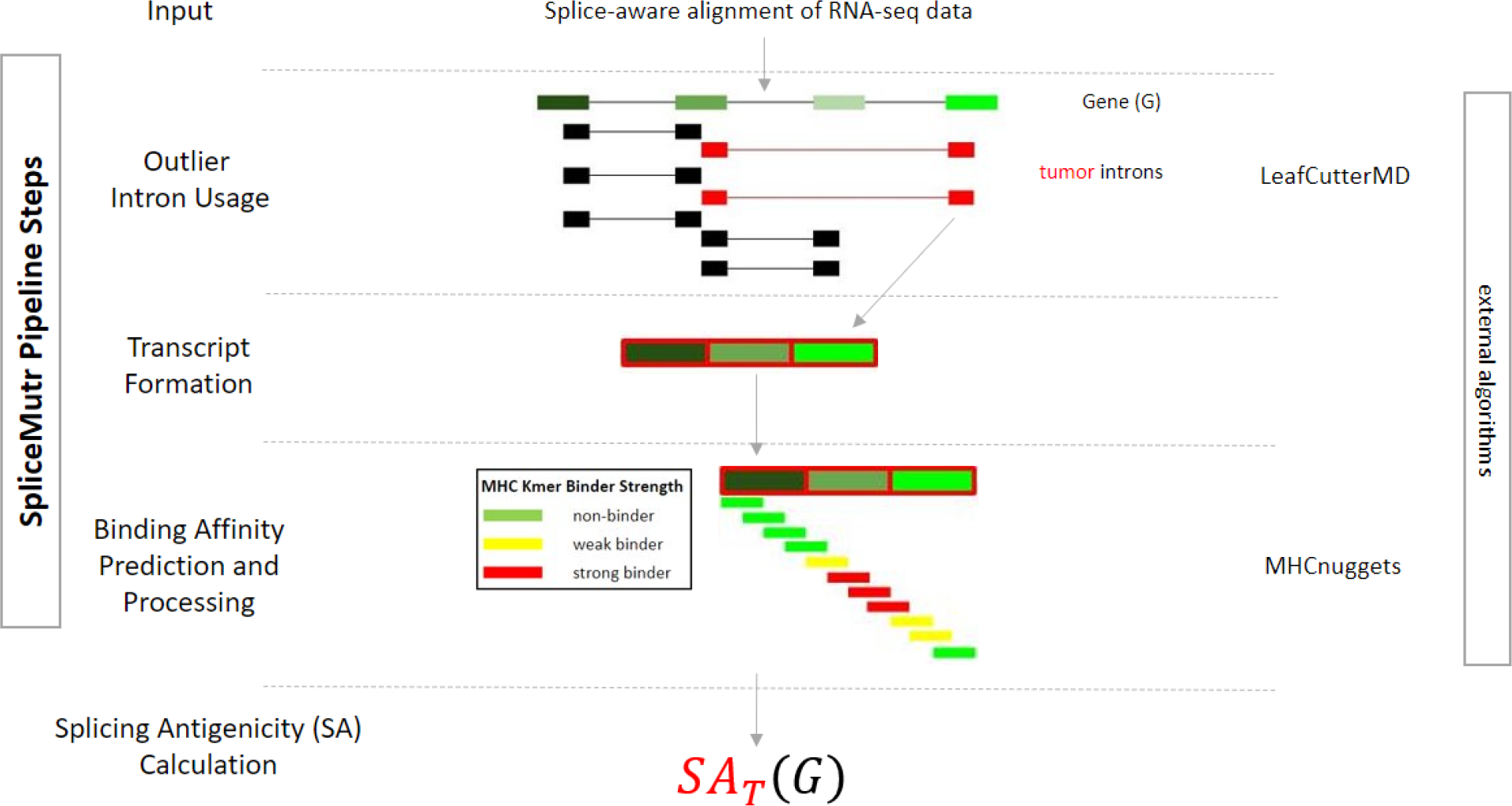
**The SpliceMutr pipeline using LeafCutterMD outlier splicing vs LeafCutter differential splicing.**

**Fig S4.**
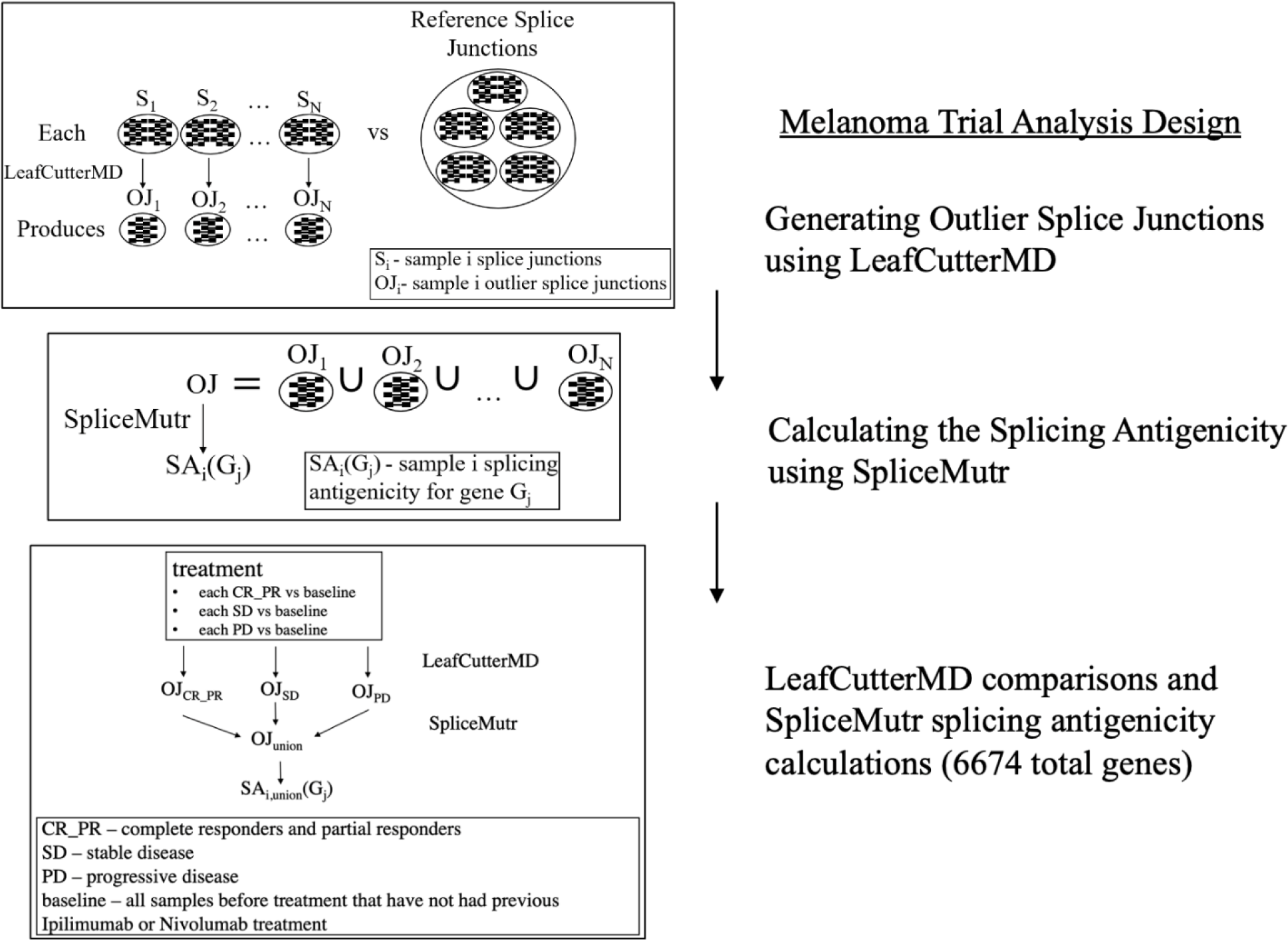
**The melanoma cohort analysis design.**

**Fig. S5.**
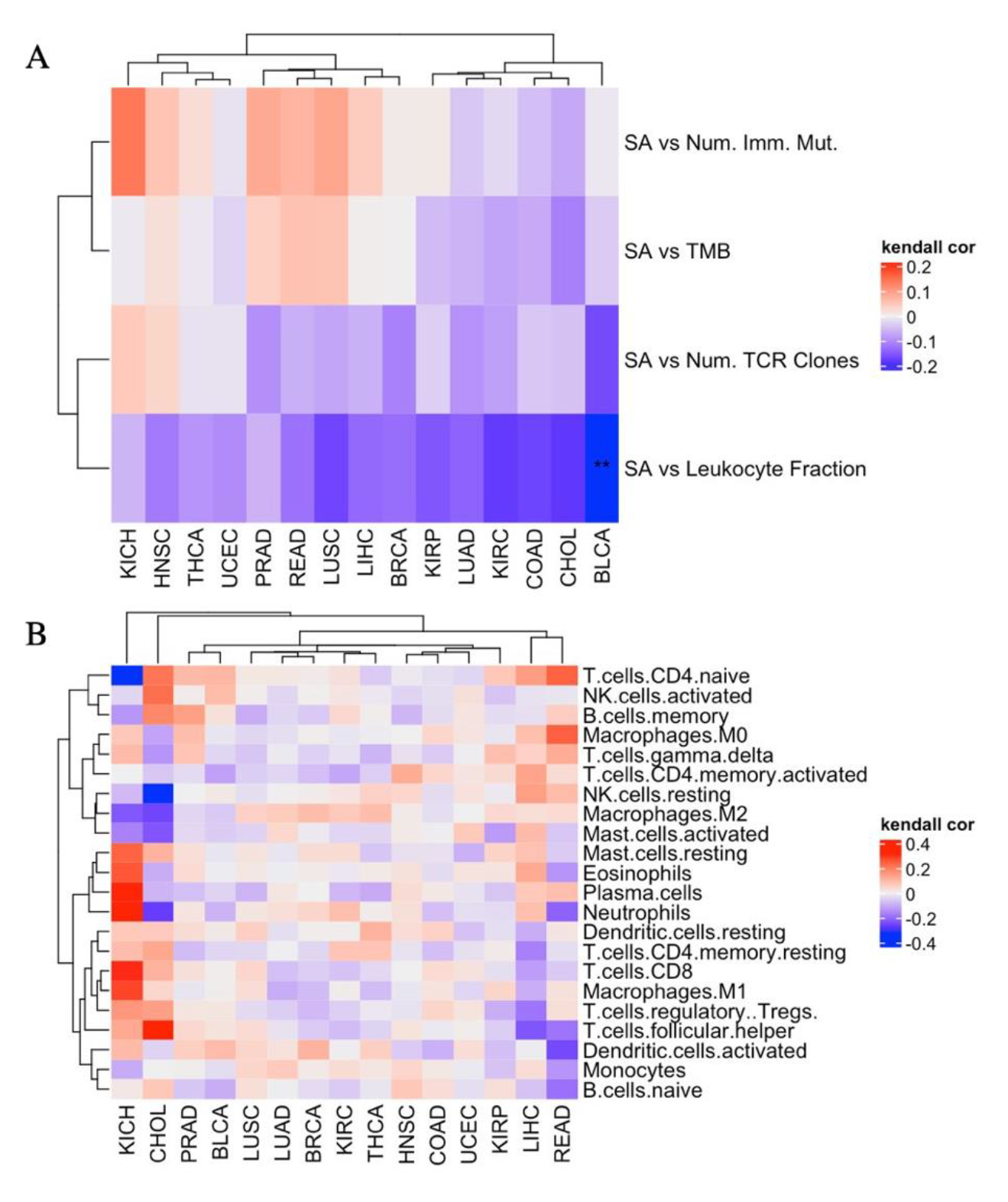
(A) The splicing antigenicity averaged across all genes per sample correlated to the TCR clonality, the number of immunogenic mutations, the leukocyte fraction, and the TMB per TCGA cancer subtype. (B) The splicing antigenicity correlated to CIBERSORT-calculated immune cell proportions per TCGA cancer subtype. (*: p-value BH < 0.05 and |tau| >= 0.1, Kendal Tau test).

**Fig. S6.**
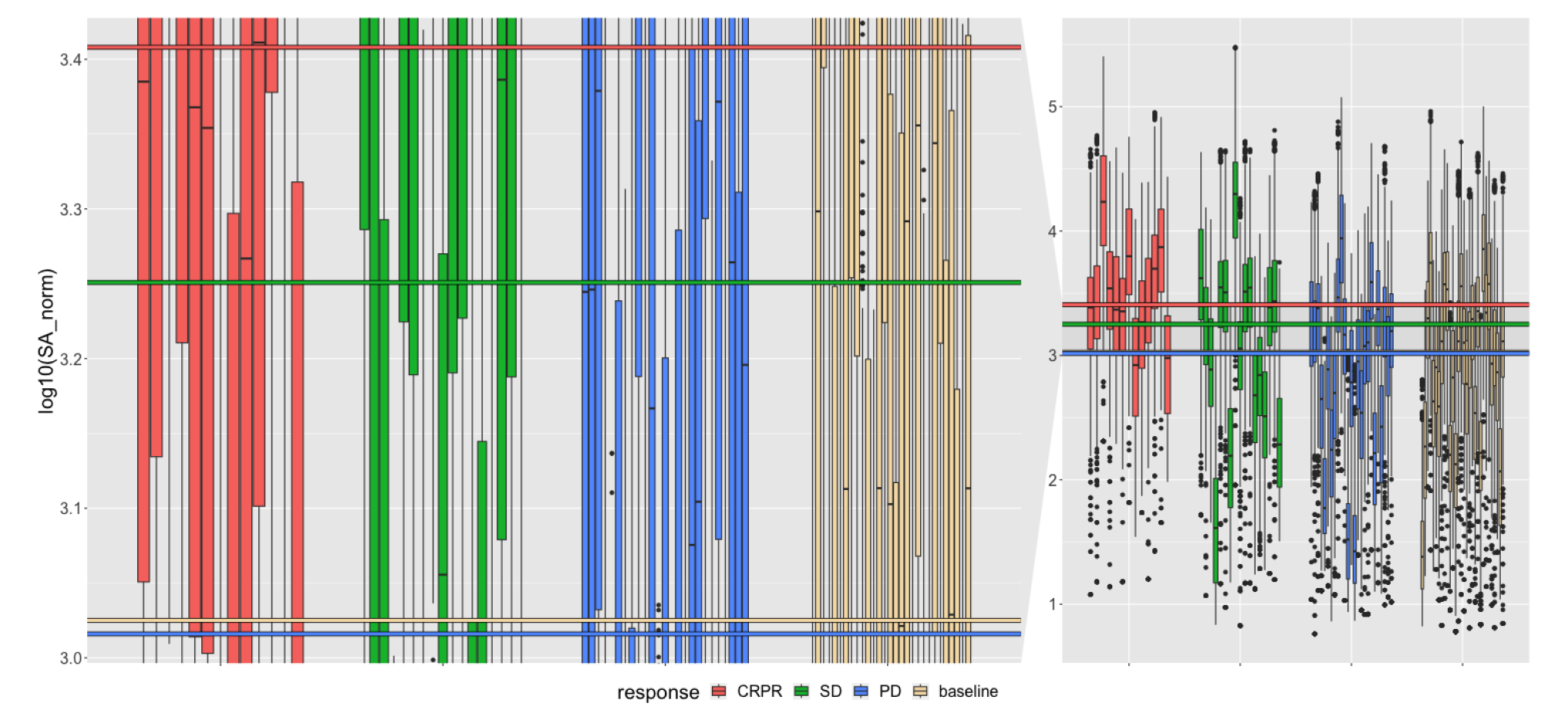
The splicing antigenicity for the subset of splice junctions derived from the top twenty genes with the highest splicing antigenicity per sample. * p-value<=0.05, ** p-value<=0.005, *** p-value<=0.0005, **** p-value<=0.00005.

## References

1. Afsari, B., Guo, T., Considine, M., Florea, L., Kagohara, L. T., Stein-O’Brien, G. L., Kelley, D., Flam, E., Zambo, K. D., Ha, P. K., Geman, D., Ochs, M. F., Califano, J. A., Gaykalova, D. A., Favorov, A. v, & Fertig, E. J. (2018). Splice Expression Variation Analysis (SEVA) for inter-tumor heterogeneity of gene isoform usage in cancer. Bioinformatics, 34(11), 1859–1867. https://doi.org/10.1093/bioinformatics/bty004

2. Aguet, F., Anand, S., Ardlie, K. G., Gabriel, S., Getz, G. A., Graubert, A., Hadley, K., Handsaker, R. E., Huang, K. H., Kashin, S., Li, X., MacArthur, D. G., Meier, S. R., Nedzel, J. L., Nguyen, D. T., Segrè, A. v., Todres, E., Balliu, B., Barbeira, A. N., … Volpi, S. (2020). The GTEx Consortium atlas of genetic regulatory effects across human tissues. Science, 369(6509), 1318–1330. https://doi.org/10.1126/science.aaz1776

3. Anagnostou, V., Bruhm, D. C., Niknafs, N., White, J. R., Shao, X. M., Sidhom, J. W., Stein, J., Tsai, H. L., Wang, H., Belcaid, Z., Murray, J., Balan, A., Ferreira, L., Ross-Macdonald, P., Wind-Rotolo, M., Baras, A. S., Taube, J., Karchin, R., Scharpf, R. B., Grasso, C., … Velculescu, V. E. (2020). Integrative Tumor and Immune Cell Multi-omic Analyses Predict Response to Immune Checkpoint Blockade in Melanoma. Cell reports. Medicine, 1(8), 100139. https://doi.org/10.1016/j.xcrm.2020.100139

4. Chen, B., Khodadoust, M. S., Liu, C. L., Newman, A. M., & Alizadeh, A. A. (2018). Profiling Tumor Infiltrating Immune Cells with CIBERSORT (pp. 243–259). https://doi.org/10.1007/978-1-4939-7493-1_12

5. Dobin, A., Davis, C. A., Schlesinger, F., Drenkow, J., Zaleski, C., Jha, S., Batut, P., Chaisson, M., & Gingeras, T. R. (2013). STAR: ultrafast universal RNA-seq aligner. Bioinformatics, 29(1), 15–21. https://doi.org/10.1093/bioinformatics/bts635

6. Eisenhauer, E. A., Therasse, P., Bogaerts, J., Schwartz, L. H., Sargent, D., Ford, R., Dancey, J., Arbuck, S., Gwyther, S., Mooney, M., Rubinstein, L., Shankar, L., Dodd, L., Kaplan, R., Lacombe, D., & Verweij, J. (2009). New response evaluation criteria in solid tumours: Revised RECIST guideline (version 1.1). European Journal of Cancer, 45(2), 228–247. https://doi.org/10.1016/j.ejca.2008.10.026

7. Grasso, C. S., Tsoi, J., Onyshchenko, M., Abril-Rodriguez, G., Ross-Macdonald, P., Wind-Rotolo, M., Champhekar, A., Medina, E., Torrejon, D. Y., Shin, D. S., Tran, P., Kim, Y. J., Puig-Saus, C., Campbell, K., Vega-Crespo, A., Quist, M., Martignier, C., Luke, J. J., Wolchok, J. D., Johnson, D. B., … Ribas, A. (2020). Conserved Interferon-γ Signaling Drives Clinical Response to Immune Checkpoint Blockade Therapy in Melanoma. Cancer cell, 38(4), 500– 515.e3. https://doi.org/10.1016/j.ccell.2020.08.005

8. Haen, S. P., Löffler, M. W., Rammensee, H.-G., & Brossart, P. (2020). Towards new horizons: characterization, classification and implications of the tumour antigenic repertoire. Nature Reviews Clinical Oncology, 17(10), 595–610. https://doi.org/10.1038/s41571-020-0387-x

9. Hong, M. M. Y., & Maleki Vareki, S. (2022). Addressing the Elephant in the Immunotherapy Room: Effector T-Cell Priming versus Depletion of Regulatory T-Cells by Anti-CTLA-4 Therapy. Cancers, 14(6), 1580. https://doi.org/10.3390/cancers14061580

10. Hu, K., Lou, L., Ye, J., & Zhang, S. (2015). Prognostic role of the neutrophil-lymphocyte ratio in renal cell carcinoma: a meta-analysis. BMJ Open, 5(4), e006404–e006404. https://doi.org/10.1136/bmjopen-2014-006404

11. Inoue, D., Chew, G.-L., Liu, B., Michel, B. C., Pangallo, J., D’Avino, A. R., Hitchman, T., North, K., Lee, S. C.-W., Bitner, L., Block, A., Moore, A. R., Yoshimi, A., Escobar-Hoyos, L., Cho, H., Penson, A., Lu, S. X., Taylor, J., Chen, Y., … Bradley, R. K. (2019). Spliceosomal disruption of the non-canonical BAF complex in cancer. Nature, 574(7778), 432–436. https://doi.org/10.1038/s41586-019-1646-9

12. Jayasinghe, R. G., Cao, S., Gao, Q., Wendl, M. C., Vo, N. S., Reynolds, S. M., Zhao, Y., Climente-González, H., Chai, S., Wang, F., Varghese, R., Huang, M., Liang, W.-W., Wyczalkowski, M. A., Sengupta, S., Li, Z., Payne, S. H., Fenyö, D., Miner, J. H., … Mariamidze, A. (2018a). Systematic Analysis of Splice-Site-Creating Mutations in Cancer. Cell Reports, 23(1), 270–281.e3. https://doi.org/10.1016/j.celrep.2018.03.052

13. Jayasinghe, R. G., Cao, S., Gao, Q., Wendl, M. C., Vo, N. S., Reynolds, S. M., Zhao, Y., Climente-González, H., Chai, S., Wang, F., Varghese, R., Huang, M., Liang, W.-W., Wyczalkowski, M. A., Sengupta, S., Li, Z., Payne, S. H., Fenyö, D., Miner, J. H., … Mariamidze, A. (2018b). Systematic Analysis of Splice-Site-Creating Mutations in Cancer. Cell Reports, 23(1), 270–281.e3. https://doi.org/10.1016/j.celrep.2018.03.052

14. Jenkinson, G., Li, Y. I., Basu, S., Cousin, M. A., Oliver, G. R., & Klee, E. W. (2020). LeafCutterMD: an algorithm for outlier splicing detection in rare diseases. Bioinformatics, 36(17), 4609–4615. https://doi.org/10.1093/bioinformatics/btaa259

15. Kahles, A., Lehmann, K.-V., Toussaint, N. C., Hüser, M., Stark, S. G., Sachsenberg, T., Stegle, O., Kohlbacher, O., Sander, C., Rätsch, G., Caesar-Johnson, S. J., Demchok, J. A., Felau, I., Kasapi, M., Ferguson, M. L., Hutter, C. M., Sofia, H. J., Tarnuzzer, R., Wang, Z., … Mariamidze, A. (2018). Comprehensive Analysis of Alternative Splicing Across Tumors from 8,705 Patients. Cancer Cell, 34(2), 211–224.e6. https://doi.org/10.1016/j.ccell.2018.07.001

16. Kahles, A., Ong, C. S., Zhong, Y., & Rätsch, G. (2016). *SplAdder*: identification, quantification and testing of alternative splicing events from RNA-Seq data. Bioinformatics, 32(12), 1840–1847. https://doi.org/10.1093/bioinformatics/btw076

17. Kouyama, Y., Masuda, T., Fujii, A., Ogawa, Y., Sato, K., Tobo, T., Wakiyama, H., Yoshikawa, Y., Noda, M., Tsuruda, Y., Kuroda, Y., Eguchi, H., Ishida, F., Kudo, S., & Mimori, K. (2019). Oncogenic splicing abnormalities induced by <*scp*>*DEAD*</*scp*> -*Box Helicase 56* amplification in colorectal cancer. Cancer Science, 110(10), 3132–3144. https://doi.org/10.1111/cas.14163

18. Lakens, Daniël. “Calculating and Reporting Effect Sizes to Facilitate Cumulative Science: A Practical Primer for T-Tests and ANOVAS.” Frontiers in Psychology, vol. 4, 2013, https://doi.org/10.3389/fpsyg.2013.00863.

19. Li, H.-D., Funk, C. C., & Price, N. D. (2020). iREAD: a tool for intron retention detection from RNA-seq data. BMC Genomics, 21(1), 128. https://doi.org/10.1186/s12864-020-6541-0

20. Li, K., Tandurella, J. A., Gai, J., Zhu, Q., Lim, S. J., Thomas, D. L., Xia, T., Mo, G., Mitchell, J. T., Montagne, J., Lyman, M., Danilova, L. v., Zimmerman, J. W., Kinny-Köster, B., Zhang, T., Chen, L., Blair, A. B., Heumann, T., Parkinson, R., … Zheng, L. (2022). Multi-omic analyses of changes in the tumor microenvironment of pancreatic adenocarcinoma following neoadjuvant treatment with anti-PD-1 therapy. Cancer Cell, 40(11), 1374–1391.e7. https://doi.org/10.1016/j.ccell.2022.10.001

21. Li, Y. I., Knowles, D. A., Humphrey, J., Barbeira, A. N., Dickinson, S. P., Im, H. K., & Pritchard, J. K. (2018). Annotation-free quantification of RNA splicing using LeafCutter. Nature Genetics, 50(1), 151–158. https://doi.org/10.1038/s41588-017-0004-9

22. Liu, Z., Yoshimi, A., Wang, J., Cho, H., Chun-Wei Lee, S., Ki, M., Bitner, L., Chu, T., Shah, H., Liu, B., Mato, A. R., Ruvolo, P., Fabbri, G., Pasqualucci, L., Abdel-Wahab, O., & Rabadan, R. (2020a). Mutations in the RNA Splicing Factor SF3B1 Promote Tumorigenesis through MYC Stabilization. Cancer Discovery, 10(6), 806–821. https://doi.org/10.1158/2159-8290.CD-19-1330

23. Liu, Z., Yoshimi, A., Wang, J., Cho, H., Chun-Wei Lee, S., Ki, M., Bitner, L., Chu, T., Shah, H., Liu, B., Mato, A. R., Ruvolo, P., Fabbri, G., Pasqualucci, L., Abdel-Wahab, O., & Rabadan, R. (2020b). Mutations in the RNA Splicing Factor SF3B1 Promote Tumorigenesis through MYC Stabilization. Cancer Discovery, 10(6), 806–821. https://doi.org/10.1158/2159-8290.CD-19-1330

24. Lu, S. X., de Neef, E., Thomas, J. D., Sabio, E., Rousseau, B., Gigoux, M., Knorr, D. A., Greenbaum, B., Elhanati, Y., Hogg, S. J., Chow, A., Ghosh, A., Xie, A., Zamarin, D., Cui, D., Erickson, C., Singer, M., Cho, H., Wang, E., … Bradley, R. K. (2021). Pharmacologic modulation of RNA splicing enhances antitumor immunity. Cell, 184(15), 4032–4047.e31. https://doi.org/10.1016/j.cell.2021.05.038

25. Maguire, S. L., Leonidou, A., Wai, P., Marchiò, C., Ng, C. K., Sapino, A., Salomon, A., Reis-Filho, J. S., Weigelt, B., & Natrajan, R. C. (2015). *SF3B1* mutations constitute a novel therapeutic target in breast cancer. The Journal of Pathology, 235(4), 571–580. https://doi.org/10.1002/path.4483

26. Middleton, R., Gao, D., Thomas, A., Singh, B., Au, A., Wong, J. J.-L., Bomane, A., Cosson, B., Eyras, E., Rasko, J. E. J., & Ritchie, W. (2017). IRFinder: assessing the impact of intron retention on mammalian gene expression. Genome Biology, 18(1), 51. https://doi.org/10.1186/s13059-017-1184-4

27. Miller, A., Asmann, Y., Cattaneo, L., Braggio, E., Keats, J., Auclair, D., Lonial, S., Russell, S. J., & Stewart, A. K. (2017). High somatic mutation and neoantigen burden are correlated with decreased progression-free survival in multiple myeloma. Blood Cancer Journal, 7(9), e612–e612. https://doi.org/10.1038/bcj.2017.94

28. Niknafs, N., Balan, A., Cherry, C., Hummelink, K., Monkhorst, K., Shao, X. M., Belcaid, Z., Marrone, K. A., Murray, J., Smith, K. N., Levy, B., Feliciano, J., Hann, C. L., Lam, V., Pardoll, D. M., Karchin, R., Seiwert, T. Y., Brahmer, J. R., Forde, P. M., Velculescu, V. E., … Anagnostou, V. (2023). Persistent mutation burden drives sustained antitumor immune responses. Nature medicine, 29(2), 440–449. https://doi.org/10.1038/s41591-022-02163-w

29. Orenbuch, R., Filip, I., Comito, D., Shaman, J., Pe’er, I., & Rabadan, R. (2020). arcasHLA: high-resolution HLA typing from RNAseq. Bioinformatics, 36(1), 33–40. https://doi.org/10.1093/bioinformatics/btz474

30. Paul, S., Weiskopf, D., Angelo, M. A., Sidney, J., Peters, B., & Sette, A. (2013). HLA Class I Alleles Are Associated with Peptide-Binding Repertoires of Different Size, Affinity, and Immunogenicity. The Journal of Immunology, 191(12), 5831–5839. https://doi.org/10.4049/jimmunol.1302101

31. Richman, L. P., Vonderheide, R. H., & Rech, A. J. (2019). Neoantigen Dissimilarity to the Self-Proteome Predicts Immunogenicity and Response to Immune Checkpoint Blockade. Cell Systems, 9(4), 375–382.e4. https://doi.org/10.1016/j.cels.2019.08.009

32. Riviere, P., Goodman, A. M., Okamura, R., Barkauskas, D. A., Whitchurch, T. J., Lee, S., Khalid, N., Collier, R., Mareboina, M., Frampton, G. M., Fabrizio, D., Sharabi, A. B., Kato, S., & Kurzrock, R. (2020). High Tumor Mutational Burden Correlates with Longer Survival in Immunotherapy-Naïve Patients with Diverse Cancers. Molecular Cancer Therapeutics, 19(10), 2139–2145. https://doi.org/10.1158/1535-7163.MCT-20-0161

33. Seidel, J. A., Otsuka, A., & Kabashima, K. (2018). Anti-PD-1 and Anti-CTLA-4 Therapies in Cancer: Mechanisms of Action, Efficacy, and Limitations. Frontiers in Oncology, 8. https://doi.org/10.3389/fonc.2018.00086

34. Seiler, M., Peng, S., Agrawal, A. A., Palacino, J., Teng, T., Zhu, P., Smith, P. G., Buonamici, S., Yu, L., Caesar-Johnson, S. J., Demchok, J. A., Felau, I., Kasapi, M., Ferguson, M. L., Hutter, C. M., Sofia, H. J., Tarnuzzer, R., Wang, Z., Yang, L., … Mariamidze, A. (2018a). Somatic Mutational Landscape of Splicing Factor Genes and Their Functional Consequences across 33 Cancer Types. Cell Reports, 23(1), 282–296.e4. https://doi.org/10.1016/j.celrep.2018.01.088

35. Shao, X. M., Bhattacharya, R., Huang, J., Sivakumar, I. K. A., Tokheim, C., Zheng, L., Hirsch, D., Kaminow, B., Omdahl, A., Bonsack, M., Riemer, A. B., Velculescu, V. E., Anagnostou, V., Pagel, K. A., & Karchin, R. (2020). High-Throughput Prediction of MHC Class I and II Neoantigens with MHCnuggets. Cancer Immunology Research, 8(3), 396–408. https://doi.org/10.1158/2326-6066.CIR-19-0464

36. Shirai, C. L., White, B. S., Tripathi, M., Tapia, R., Ley, J. N., Ndonwi, M., Kim, S., Shao, J., Carver, A., Saez, B., Fulton, R. S., Fronick, C., O’Laughlin, M., Lagisetti, C., Webb, T. R., Graubert, T. A., & Walter, M. J. (2017). Mutant U2AF1-expressing cells are sensitive to pharmacological modulation of the spliceosome. Nature Communications, 8(1), 14060. https://doi.org/10.1038/ncomms14060

37. Shuai, S., Suzuki, H., Diaz-Navarro, A., Nadeu, F., Kumar, S. A., Gutierrez-Fernandez, A., Delgado, J., Pinyol, M., López-Otín, C., Puente, X. S., Taylor, M. D., Campo, E., & Stein, L. D. (2019). The U1 spliceosomal RNA is recurrently mutated in multiple cancers. Nature, 574(7780), 712–716. https://doi.org/10.1038/s41586-019-1651-z

38. Sidney, J., Southwood, S., Moore, C., Oseroff, C., Pinilla, C., Grey, H. M., & Sette, A. (2013). Measurement of MHC/Peptide Interactions by Gel Filtration or Monoclonal Antibody Capture. Current Protocols in Immunology, 100(1). https://doi.org/10.1002/0471142735.im1803s100

39. Szolek, A., Schubert, B., Mohr, C., Sturm, M., Feldhahn, M., & Kohlbacher, O. (2014). OptiType: precision HLA typing from next-generation sequencing data. Bioinformatics, 30(23), 3310–3316. https://doi.org/10.1093/bioinformatics/btu548

40. Thorsson, V., Gibbs, D. L., Brown, S. D., Wolf, D., Bortone, D. S., Ou Yang, T.-H., Porta-Pardo, E., Gao, G. F., Plaisier, C. L., Eddy, J. A., Ziv, E., Culhane, A. C., Paull, E. O., Sivakumar, I. K. A., Gentles, A. J., Malhotra, R., Farshidfar, F., Colaprico, A., Parker, J. S., … Mariamidze, A. (2018). The Immune Landscape of Cancer. Immunity, 48(4), 812–830.e14. https://doi.org/10.1016/j.immuni.2018.03.023

41. Trincado, J. L., Reixachs-Solé, M., Pérez-Granado, J., Fugmann, T., Sanz, F., Yokota, J., & Eyras, E. (2021). ISOTOPE: ISOform-guided prediction of epiTOPEs in cancer. PLOS Computational Biology, 17(9), e1009411. https://doi.org/10.1371/journal.pcbi.1009411

42. Wang, G., Yin, H., Li, B., Yu, C., Wang, F., Xu, X., Cao, J., Bao, Y., Wang, L., Abbasi, A. A., Bajic, V. B., Ma, L., & Zhang, Z. (2019). Characterization and identification of long non-coding RNAs based on feature relationship. Bioinformatics, 35(17), 2949–2956. https://doi.org/10.1093/bioinformatics/btz008

43. Wilks, C., Zheng, S. C., Chen, F. Y., Charles, R., Solomon, B., Ling, J. P., Imada, E. L., Zhang, D., Joseph, L., Leek, J. T., Jaffe, A. E., Nellore, A., Collado-Torres, L., Hansen, K. D., & Langmead, B. (2021). recount3: summaries and queries for large-scale RNA-seq expression and splicing. Genome Biology, 22(1), 323. https://doi.org/10.1186/s13059-021-02533-6

44. Wu, M., Huang, Q., Xie, Y., Wu, X., Ma, H., Zhang, Y., & Xia, Y. (2022). Improvement of the anticancer efficacy of PD-1/PD-L1 blockade via combination therapy and PD-L1 regulation. Journal of Hematology & Oncology, 15(1), 24. https://doi.org/10.1186/s13045-022-01242-2

45. Xu, Z., Dai, J., Wang, D., Lu, H., Dai, H., Ye, H., Gu, J., Chen, S., & Huang, B. (2019). <p>Assessment of tumor mutation burden calculation from gene panel sequencing data</p>. OncoTargets and Therapy, *Volume* 12, 3401–3409. https://doi.org/10.2147/OTT.S196638

46. Yoshihara, Kosuke, et al. “Inferring Tumour Purity and Stromal and Immune Cell Admixture from Expression Data.” Nature Communications, vol. 4, no. 1, 2013.

47. Zhang, J., Ali, A. M., Lieu, Y. K., Liu, Z., Gao, J., Rabadan, R., Raza, A., Mukherjee, S., & Manley, J. L. (2019). Disease-Causing Mutations in SF3B1 Alter Splicing by Disrupting Interaction with SUGP1. Molecular Cell, 76(1), 82–95.e7. https://doi.org/10.1016/j.molcel.2019.07.017

