## Supplementary Figures for "SpliceMutr enables pan-cancer analysis of splicing-derived neoantigen burden in tumors"

### Supplement

| Study Abbreviation | Study Name |
|---|---|
| LAML | Acute Myeloid Leukemia |
| ACC | Adrenocortical carcinoma |
| BLCA | Bladder Urothelial Carcinoma |
| LGG | Brain Lower Grade Glioma |
| BRCA | Breast invasive carcinoma |
| CESC | Cervical squamous cell carcinoma and endocervical |
| CHOL | Cholangiocarcinoma |
| LCML | Chronic Myelogenous Leukemia |
| COAD | Colon adenocarcinoma |
| ESCA | Esophageal carcinoma |
| GBM | Glioblastoma multiforme |
| HNSC | Head and Neck squamous cell carcinoma |
| KICH | Kidney Chromophobe |
| KIRC | Kidney renal clear cell carcinoma |
| KIRP | Kidney renal papillary cell carcinoma |
| LIHC | Liver hepatocellular carcinoma |
| LUAD | Lung adenocarcinoma |
| LUSC | Lung squamous cell carcinoma |
| DLBC | Lymphoid Neoplasm Diffuse Large B-cell Lymphoma |
| MESO | Mesothelioma |
| MISC | Miscellaneous |
| OV | Ovarian serous cystadenocarcinoma |
| PAAD | Pancreatic adenocarcinoma |
| PCPG | Pheochromocytoma and Paraganglioma |
| PRAD | Prostate adenocarcinoma |
| READ | Rectum adenocarcinoma |
| SARC | Sarcoma |
| SKCM | Skin Cutaneous Melanoma |
| STAD | Stomach adenocarcinoma |
| TGCT | Testicular Germ Cell Tumors |
| THYM | Thymoma |
| THCA | Thyroid carcinoma |
| UCS | Uterine Carcinosarcoma |
| UCEC | Uterine Corpus Endometrial Carcinoma |
| UVM | Uveal Melanoma |

**Table S1. TCGA cancer study abbreviations and names.**

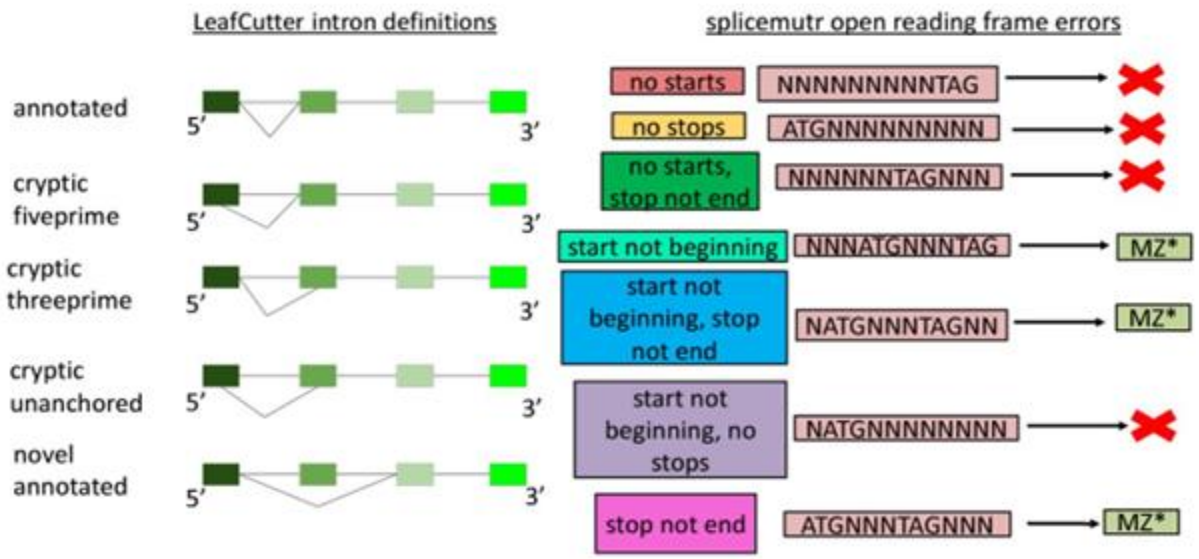


**Fig S1. LeafCutter intron definitions and SpliceMutr open reading frame errors.** The LeafCutter intron definitions are a visual representation of the LeafCutter intron definitions with respect to a mock gene model. Annotated introns have a pair of splice sites located at the genomic coordinates of a documented intron. Cryptic intron types have either one or both splice sites located at genomic coordinates not associated with a documented intron. Novel annotated introns have a pair of splice sites individually located at the genomic coordinates of documented introns but not at splice site locations documented to be joined together. The splicemutr open reading frame errors are a visual representation of the open reading frame (ORF) errors that splicemutr documents during transcript formation. The full range of the DNA equivalent of stop codons are defined as such during transcript formation, but this example only features the TAG stop codon. If an open reading frame can be found after modification of a reference transcript by a differentially used intron, then the transcript is translated with the appropriate error documented. If the modified transcript has unchanged ORF start and end codons relative to the reference, then the modified transcript is documented as being immediately translatable.

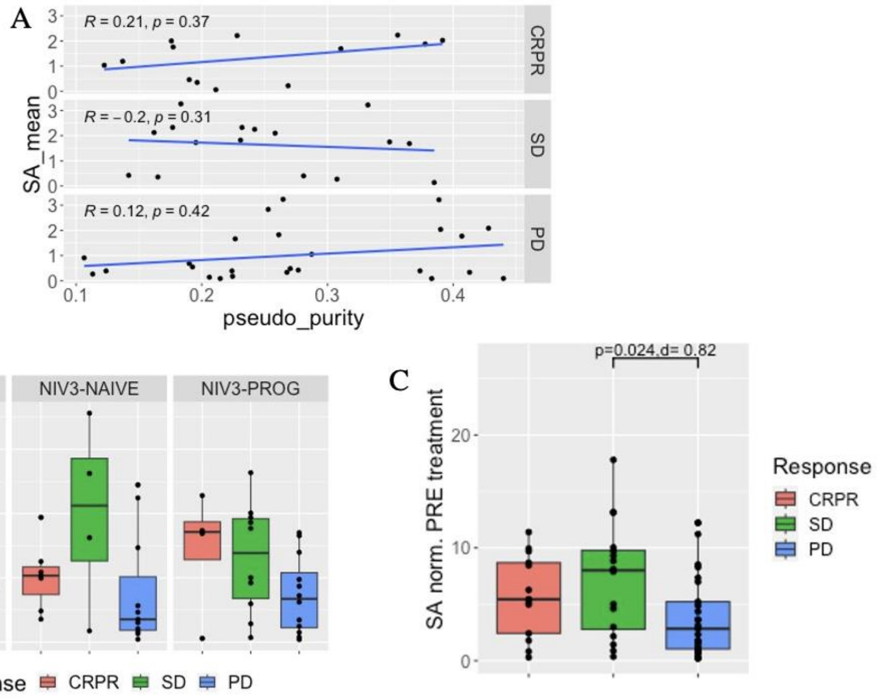


**Fig. S2. Per-patient splicing antigenicity of ICI-treated melanoma patients per treatment arm and response type before treatment.** (A) The per-patient pseudo purity compared to the mean splicing antigenicity averaged across genes per-response type for all treatment arms. Kendall Tau test.(B) The mean splicing antigenicity averaged across genes per patient and normalized by the pseudo purity, for each treatment arm. Wilcoxon test with false discovery rate adjustment and Cohens d. (C) The mean splicing antigenicity averaged across genes per patient and normalized by the pseudo purity, for all treatment arms combined. Wilcoxon test and Cohens d.

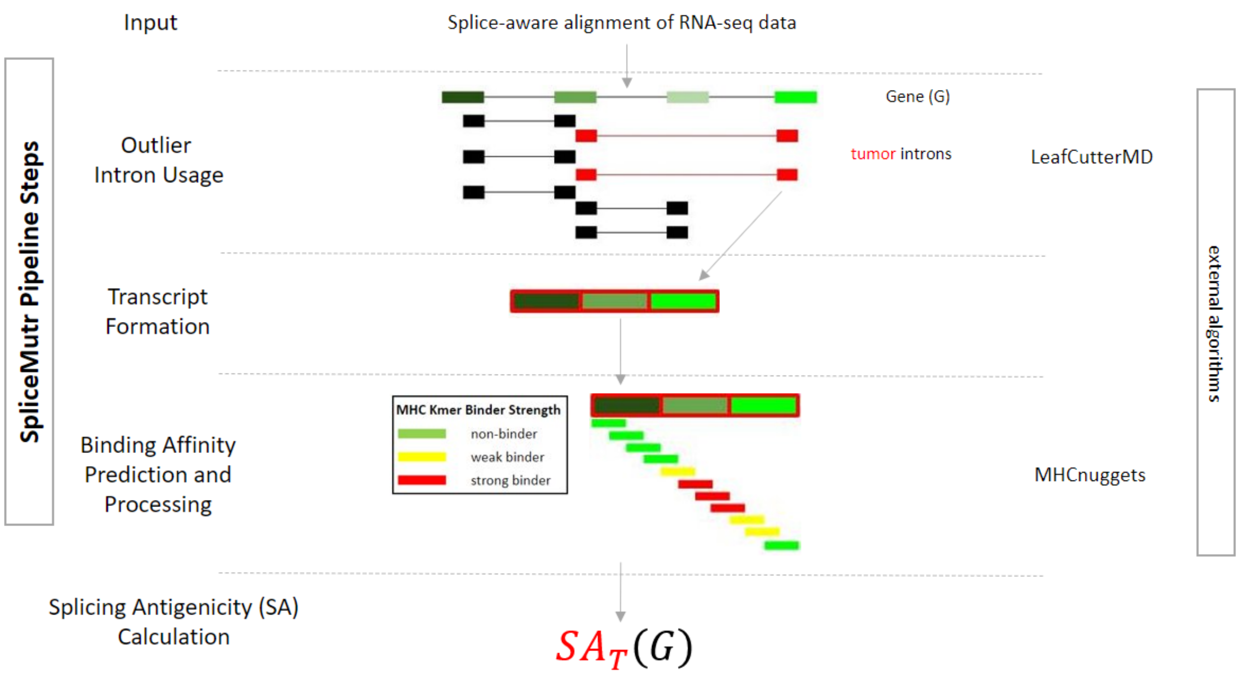


**Fig. S3. The SpliceMutr pipeline using LeafCutterMD outlier splicing vs LeafCutter differential splicing.**

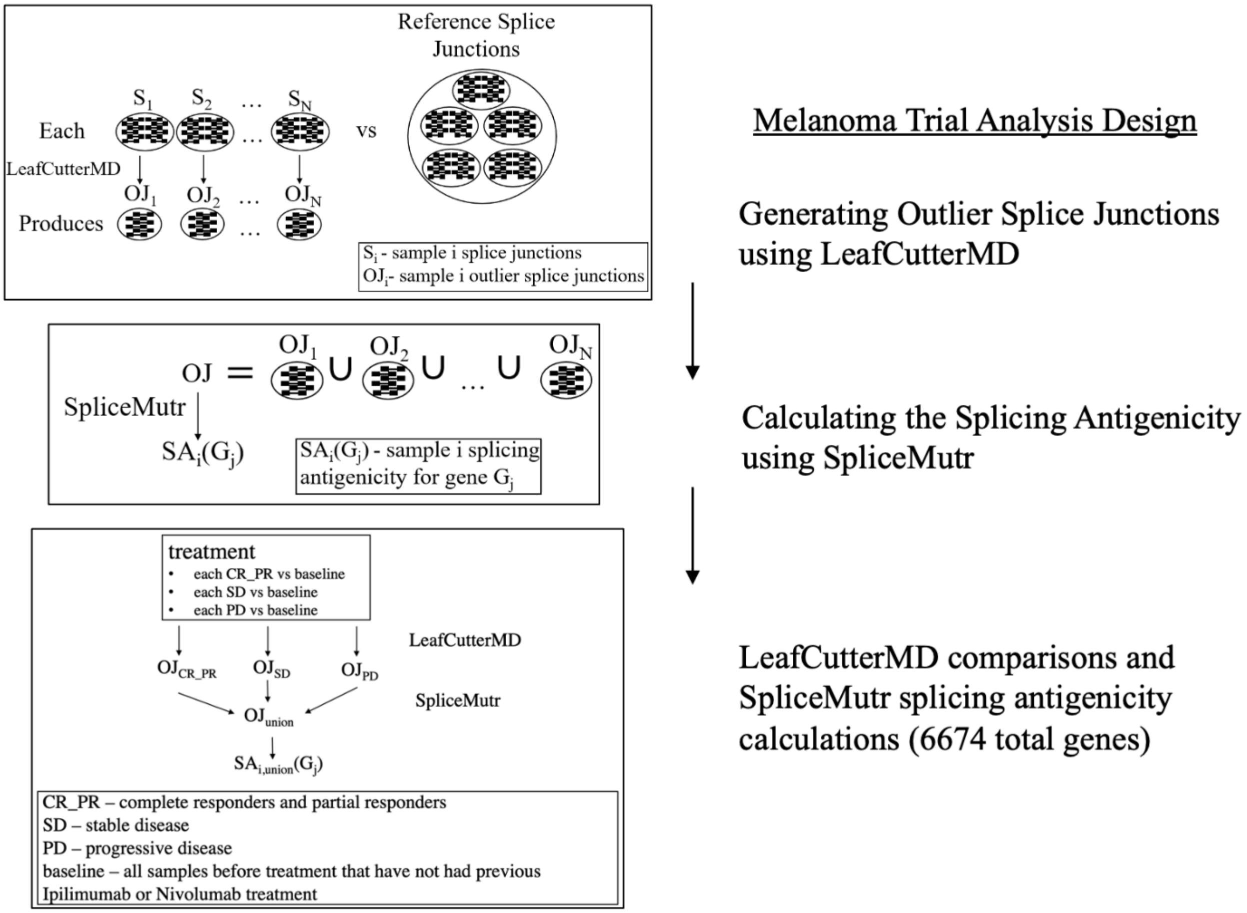

**Fig S4 The melanoma cohort analysis design.**

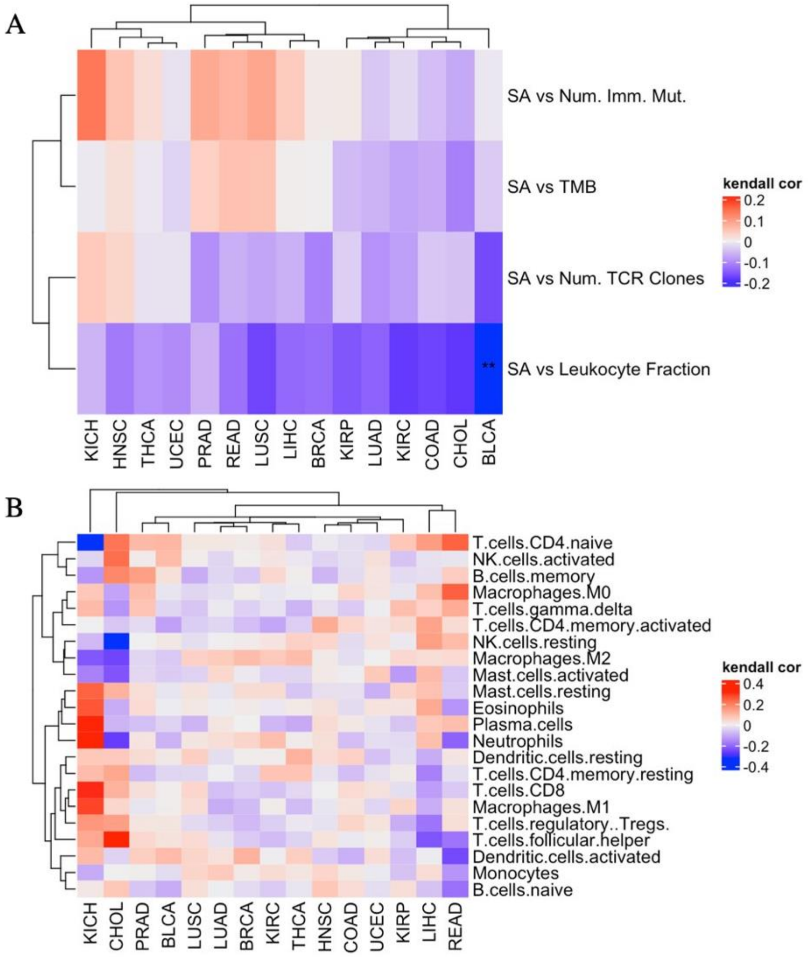


e

**Fig. S5.** (A) The splicing antigenicity averaged across all genes per sample correlated to the TCR clonality, the number of immunogenic mutations, the leukocyte fraction, and the TMB per TCGA cancer subtype. (B) The splicing antigenicity correlated to CIBERSORT-calculated immune cell proportions per TCGA cancer subtype. (*: p-value BH < 0.05 and |tau| >= 0.1, Kendal Tau test).

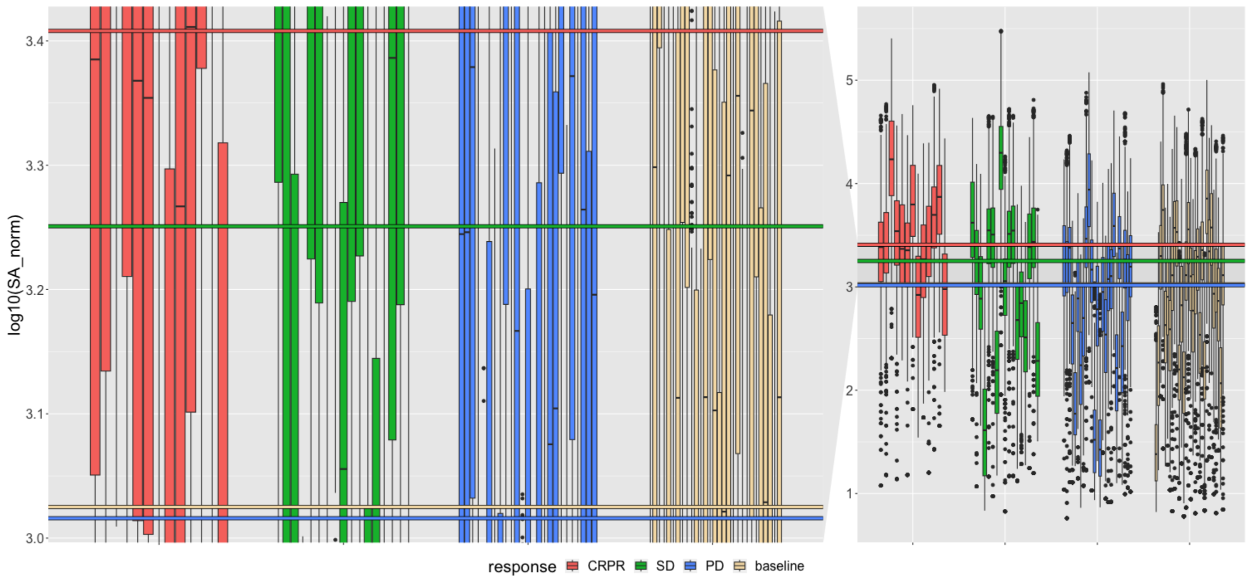


**Fig. S6.** The splicing antigenicity for the subset of splice junctions derived from the top twenty genes with the highest splicing antigenicity per sample. * p-value<=0.05, ** p-value<=0.005, *** p-value<=0.0005, **** p-value<=0.00005.
